# I-SVVS: Integrative stochastic variational variable selection to explore joint patterns of multi-omics microbiome data

**DOI:** 10.1101/2023.08.18.553796

**Authors:** Tung Dang, Yushiro Fuji, Kie Kumaishi, Erika Usui, Shungo Kobori, Takumi Sato, Yusuke Toda, Kengo Sakurai, Yuji Yamasaki, Hisashi Tsujimoto, Masami Yokota Hirai, Yasunori Ichihashi, Hiroyoshi Iwata

## Abstract

High-dimensional multi-omics microbiome data plays an important role in elucidating microbial communities’ interactions with their hosts and environment in critical diseases and ecological changes. Although Bayesian clustering methods have recently been used for the integrated analysis of multi-omics data, no method designed to analyze multi-omics microbiome data has been proposed. In this study, we propose a novel framework called integrative stochastic variational variable selection (I-SVVS), which is an extension of stochastic variational variable selection for high-dimensional microbiome data. The I-SVVS approach addresses a specific Bayesian mixture model for each type of omics data, such as an infinite Dirichlet multinomial mixture model for microbiome data and an infinite Gaussian mixture model for metabolomic data. This approach is expected to reduce the computational time of the clustering process and improve the accuracy of the clustering results. Additionally, I-SVVS identifies a critical set of representative variables in multi-omics microbiome data. Three datasets from soybean, mice, and humans (each set integrated microbiome and metabolome) were used to demonstrate the potential of I-SVVS. The results indicate that I-SVVS achieved improved accuracy and faster computation compared to existing methods across all test datasets. It effectively identified key microbiome species and metabolites characterizing each cluster. For instance, the computational analysis of soybean dataset, including 377 samples with 16,943 microbiome species and 265 metabolome features, was completed in 2.18 hours using I-SVVS, compared to 2.35 days with Clusternomics and 1.12 days with iClusterPlus. The software for this analysis, written in Python, is freely available at https://github.com/tungtokyo1108/I-SVVS.

## Introduction

Owing to the substantial development of high-throughput technologies, high-dimensional omics data have been generated in various areas, such as agriculture and medicine. For example, in agricultural crops, multi-omics datasets, including soil metabolites, minerals, and microbes, provide an opportunity to jointly analyze datasets to elucidate the network structure of an agroecosystem [1, 2, 3]. In medicine, the joint analysis of a multi-omics microbiome dataset plays an important role in investigating the influence of host genes and microbiome associations [4, 5, 6, 7, 8, 9, 10] or host metabolism and microbiome associations [11, 12, 13, 14] on human health and diseases.

The typical challenges in the development of computational methods for conducting joint analysis of multi-omics datasets are the problems of high dimensionality, sparsity, and multicollinearity to identify biologically meaningful associations among a large number of heterogeneous biological variables in different types of omics. Recently, several integrative approaches have been developed for the joint analysis of multiomics datasets. For example, Bayesian Consensus Clustering (BCC) [15] is a Bayesian approach that simultaneously estimates clustering specific to each omics dataset and consensus clustering by integrating all datasets. BCC introduces parameters that adjust the differences in cluster assignments between datasets, allowing for heterogeneity across different omics datasets. A consensus clustering solution was then estimated by integrating the posterior distributions of the latent cluster variables across all the datasets. However, BCC is not suitable for very large and complex datasets, because the computational complexity and memory requirements can become prohibitively high. Furthermore, the important assumption of BCC that the observed omics variables follow normal distributions limits the applicability of this method. As a more recent example, Clusternomics [16] is a probabilistic framework with hierarchical Dirichlet mixture models that rely on the existence of a consistent clustering structure across heterogeneous datasets. A context-specific cluster was developed for specific omics data to describe particular aspects of biological processes. Global clustering results from a combination of context-specific cluster assignments. Because of the two-level cluster assignment, the number of clusters in the local or global structures can be flexibly changed. However, the biological interpretability of Clusternomics is limited because many features in all omics datasets are involved in the analysis processes. The computational burden of the Clusternomics is also prohibitive. As another example, the iCluster algorithm [17] is a dimensionality reduction approach that uses a Gaussian latent variable model to infer clusters. The LASSO penalty was proposed to identify genomic variables that play important roles in the latent process. The iCluster permits the joint modeling of discrete and continuous variables. However, iCluster assumes that different omics datasets are generated from the same biological samples, which may not always be the case. Additionally, the selection of the number of clusters and penalty parameters can significantly influence the clustering solution and biological interpretability.

Currently, many accessible multi-omics microbiome datasets are becoming available [18], and it will become increasingly common to study the microbiome in relation to other omics, such as the expression of host genes and host metabolism. By integrating microbiome data (16S rRNA sequencing) with multiple sources of omics data, we may be able to elucidate the nature of host-microbe interactions, which may ultimately lead to novel discoveries. However, most current approaches are designed for the multi-omics datasets that include mRNA, microRNAs, DNA methylation, and proteomics, such as The Cancer Genome Atlas (TCGA) [19]. The major challenges associated with microbiome datasets have not yet been fully addressed in the development of integrative frameworks. For example, given the complex nature of metagenomic data, the current BCC and Clusternomics approaches cannot cluster communities into groups with similar compositions. In contrast, the Dirichlet multinomial mixture (DMM) [20, 21] is a successful method for probabilistic modeling of microbial metagenomics data. In a previous study [21], we proposed an improved framework of the DMM model called stochastic variational variable selection (SVVS). SVVS was used to predict the number of clusters and quickly identify the core set of important microbial species that contribute to the variation in different community compositions. However, combining the DMM with other mixture models for the joint analysis of a multi-omics microbiome dataset is currently one of the most challenging questions in computational biology. Each Bayesian mixture model has its own set of parameters, such as the number of clusters and prior probabilities over the cluster parameters, which can vary widely depending on the omics data. Another key challenge of Bayesian methods is that the number of biological variables becomes very large when microbial metagenomics is integrated with other omics data such as metabolism or host genomics data. Therefore, identifying the small number of representative variables that significantly contribute to the joint analysis of multi-omics microbiome datasets is crucial. Markov chain Monte Carlo (MCMC) methods, such as Gibbs sampling, which are used in BCC and Clusternomics, however, are difficult to use, given the dimensionality of the microbiome multi-omics dataset.

To address these challenges, we propose a significantly enhanced SVVS method, called integrative stochastic variational variable selection (I-SVVS). I-SVVS identifies heterogeneous patterns of sample-to-sample variability by integrating different types of datasets: count (microbiome) and continuous (metabolome). A key aspect of this method is the use of the hierarchical Dirichlet process (HDP) approach to [22] model the relationship between clusters across microbiome and metabolome datasets. This is achieved by introducing a set of shared clusters that are present in all data types. These shared clusters were modeled using the global structure of HDP. In the local structure, HDP defines a separate Dirichlet process for each cluster. This allows the distribution of data within each cluster to be modeled separately from the distribution of data across clusters. HDP has been successfully used in a wide variety of applications to analyze large datasets related to population genetics [23], protein homology detection [24, 25] and single-cell data clustering [26, 27, 28]. For the microbiome, I-SVVS uses a modeling strategy similar to that used in our previous study, that is, SVVS. For the metabolome, I-SVVS employs an infinite Gaussian mixture model (GMM) that uses a stick-breaking representation to treat the total number of clusters as a free parameter. An indicator variable was integrated into the framework of the infinite GMM to select significant metabolomic features. I-SVVS can be used for disparate analysis tasks, including joint clustering, data integration, and the identification of significant features in multi-omics data. To highlight this functionality, we applied the I-SVVS method to an integrative dataset of the microbiome and metabolome collected in our soybean experiments as well as public datasets of mice [29] and human gut disease [30].

## Materials and methods

### The integrative framework of infinite mixture models for microbiome multi-omics data

The proposed approach integrates a diverse range of data types, including metabolomics, microbiomics, ionomics, genomics, and so on. First, we introduce the following notation: *X*_*mij*_ denotes the omics variable associated with the *j′*^*th*^ (*j′* ∈ [1, …, *D*_*m*_]) omics feature in the *i*^*th*^ (*i* ∈ [1, …, *N*]) sample of the *m*^*th*^ (*m* ∈ [1, …, *M*]) omics data type. For example, an omics feature can be either a metabolite profile or a microbial species, depending on the data type. The binary latent variable *W*_*ik*_ ∈ (0, 1) denotes an indicator variable that assigns samples to the *k*^*th*^ (*k* ∈ [1, 2, …]) global-level cluster. In the global-level construction, we applied the conventional stick-breaking representation as follows:

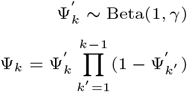

where the random variables Ψ_*k*_ denote the stick-breaking weights that satisfy 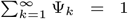 and are generated by breaking a unit-length stick into an infinite number of pieces. The indicator variable ***W*** = (*W*_*i*1_, *W*_*i*2_, …) is distributed according to **Ψ** = (Ψ_1_, Ψ_2_, …) in the following form:

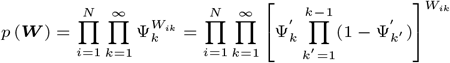

Subsequently, the indicator variable *Z*_*mkt*_ ∈ (0, 1) assigns the *k*^*th*^ global cluster to the *t*^*th*^ local cluster in the *m*^*th*^ omics data type. Similarly, we used the stick-breaking representation to construct each local-level Dirichlet process for specific omics data types as follows:

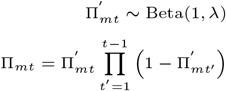

where Π_*mt*_ denotes a set of stick-breaking weights that satisfy 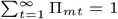 The indicator variable ***Z*** = (*Z*_*mk*1_, *Z*_*mk*2_, …) is distributed according to **Π** = (Π_*m*1_, Π_*m*2_, …) in the form

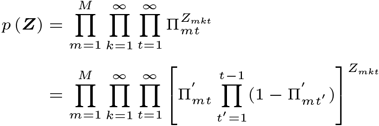

Moreover, microbiome multi-omics data sets typically include a large number of features. However, in practice, not all omics features are significant, and a large number of them may be irrelevant and negatively influence the performance of the clustering processes. Therefore, a feature selection approach is necessary to select the best omics feature subsets. We propose a binary latent variable Φ_*mij*_ which represents the feature relevance indicator. Specifically, Φ_*mij*_ = 1 means that the *j*^*th*^ feature of the *m*^*th*^ omics data type is important, otherwise, the feature *X*_*mij*_ is irrelevant.

The corresponding likelihood function of the proposed model for samples **X** = (*X*_1*ij*_, *X*_2*ij*_, …, *X*_*Mij*_) can be written as

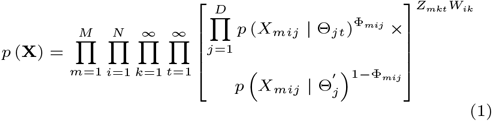

where *p* (*X*_*mij*_) is the probability density selected for the specific omis data type. Therefore, the different choices for *p* (*X*_*mij*_) allow the modeling of different types of omics data.

Here, we describe our modeling approach for specific omics datasets. In a simple case, there are two types of omics dataset (M = 2). One is microbial metagenomic data and the other is metabolite data. If *X*_1*ij*_ denotes the microbial metagenomic variable associated with the *j*^*th*^ (*j* ∈ [1, …, *D*]) taxonomic units (or species) in the *i*^*th*^ (*i* ∈ [1, …, *N*]) sample, because microbial data are the count data type, we consider the infinite Dirichlet multinomial mixture (DMM) model in our previous study [21] as follow

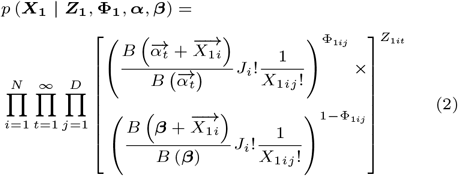

where *Z*_1*it*_ ∈ [0, 1] is an allocation variable, such that *Z*_1*it*_ = 1 if 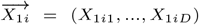 belongs to the *t* ^*th*^cluster and 0, otherwise. Φ_1*ij*_ is an indicator variable, such that Φ_1*ij*_ = 1 indicates that the *j*^*th*^ taxonomic units of *i*^*th*^ sample are important in the *t*^*th*^ cluster and follow a Dirichlet-multinomial distribution with ***α*** parameter and Φ_1*ij*_′ = 0 denotes that the *j′*^*th*^ taxonomic units of *i*^*th*^ sample is unimportant in the *t*^*th*^ cluster and follows a Dirichlet-multinomial distribution with ***β*** parameter. Function B is a m**u**ltinomia Beta function 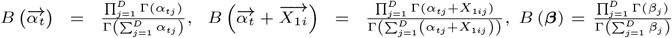 and 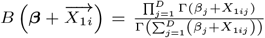. The total number of counts (i.e., sequence reads) from the *i*^*th*^ community sample was 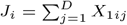.

If *X*_2*ij*_*′* is a continuous variable, we assume that it follows a normal distribution, and consider a Dirichlet process mixture model. In this study, the metabolite profile data were continuous variables. *X*_2*ij*_*′* denotes the metabolite profile variable associated with the *j′* ^*th*^ (*j′* ∈ [1, …, *D′*] metabolite profile features in the *i*^*th*^ sample. The likelihood function of the proposed model is expressed as follows:

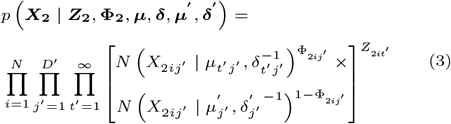

where *Z*_2*it*_*′* ∈ [0, 1] is an allocation variable such that *Z*_2*it*_*′* = 1 if 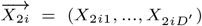 belongs to the *t ‘*^*th*^cluster and 0, otherwise. Φ_2*ij*_*′* is an indicator variable, such that Φ_2*ij*_*′* = 1 indicates that the *j′* ^*th*^ metabolite profile feature of *i*^*th*^ sample is important in the *t′* ^*th*^ cluster and follows a normal distribution with ***µ′, δ′*** parameter and Φ_2*ij*_*′* = 0 denotes that the *j′* ^*th*^ metabolite profile feature of *i*^*th*^ sample is unimportant in the *t′* ^*th*^ cluster and follows a normal distribution with ***µ’, δ’′*** parameter.

We substitute Equations 2 and 3 into Equation 1. If integration of microbiome data with metabolomics data, the likelihood function for the samples ***X*** = (*X*_1*ij*_, *X*_2*ij*_*′)* can be written as follows:

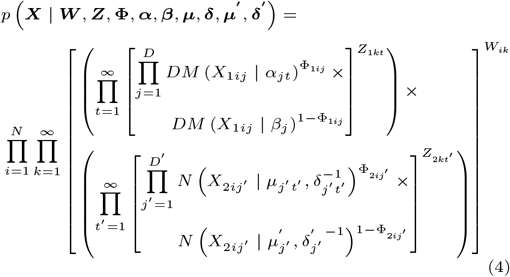

where *W*_*ik*_ ∈ [0, 1] is an allocation variable of the global cluster, such that *W*_*ik*_ = 1 if the *i*^*th*^ sample of microbiomics and metabolomics data belongs to the *k*^*th*^ global cluster and 0, otherwise. *Z*_1*kt*_ ∈ [0, 1] is an allocation variable of the local cluster for microbiome data, such as *Z*_1*kt*_ = 1 the *k*^*th*^ global cluster belongs to the *t*^*th*^ local cluster of microbiome data and 0, otherwise. *Z*_2*kt*_*′* ∈ [0, 1] is an allocation variable of a local cluster for metabolomics data, such that *Z*_2*kt*_*′* = 1 the *k* ^*th*^ global cluster belongs to the *t′*^*th*^ local cluster of metabolomics data and 0, otherwise. *DM* () denotes the Dirichlet-multinomial distribution and *N* () denotes the normal distribution. The prior distributions, that were considered specifically for all variables and parameters, are explained in detail in the Supplementary Material.

### The integrative stochastic variational variable selection (I-SVVS) approach for microbiome multi-omics data

Here, we propose an extension of the stochastic variational inference (SVI) approach, which was proposed to estimate the parameters of the infinite DMM model in our previous study [21], to learn the integrative framework of the proposed model. Given the observed omics datasets ***X***, the proposed model has a set of parameters (**Ξ**), an allocation variable of global cluster (***W***) with the unit length of the stick of the stick-breaking representation (**Ψ*′*)**, an allocation variable of the local cluster for each specific omics data (***Z***) with the unit length of the stick (**Π*′)***, the indicator variable of omics feature selection (**Φ**) and parameters (**Θ**) of the distribution *p* (***X*** | **Θ**). For example, in the case of microbiome and metabolomics data, (**Θ**) includes the parameters (***α, β***) of the Dirichlet-multinomial distributions and the parameters (***µ, δ***) of normal distributions. We then define the variational distribution of the parameters *q* (**Ξ**). In this study, we adopt the factorization assumption of mean-field variational inference, which allows for independence among the variables of the variational distribution *q* (**Ξ**). Furthermore, the proposed model integrated integrates several infinite mixture models by proposing an allocation variable for global cluster (***W***). Thus, to obtain feasible computations, truncated stick-breaking representations are considered for the global cluster at the largest value K_max_ and local cluster at the largest value T_max_. The truncation levels of the global and local clusters become variational parameters that can be automatically optimized by extending the SVI approach. The variational distribution *q* (**Ξ**) can be specifically factorized into the disjoint tractable distributions as follows:

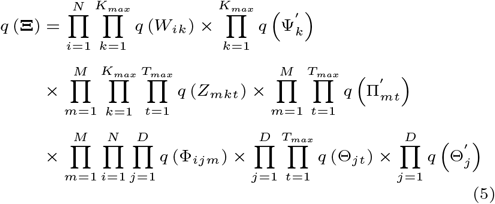

The distributions of the exponential families were selected for the variational distributions to guarantee a feasible computation of the expectations. The specific considerations for specific omics data, such as microbiome and metabolomics, are explained in detail in the Supplementary Material.

Then, the Kullback-Leibler (KL) divergence is used to evaluate the distance between the true intractable posterior distributions *p* (**Ξ** | ***X***) and *q* (**Ξ**). In previous studies, we showed that the fact indices for the computation of KL divergence were difficult. Thus, the variational framework maximizes the Evidence Lower Bound (ELBO), which equals the minimization of KL divergence, to approximate the true posterior distribution *p* (**Ξ** | ***X***). The ELBO function of the proposed method is expressed as follows:

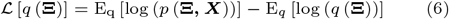

In the framework of the SVI approach, it is important to divide the variational parameters into two subgroups: local variables **Ξ**_***l***_ and global variables **Ξ**_***g***_. Expect for the global cluster variable ***W***, the numbers of local and global variables depend on the number of omics datasets used for integrated analysis. The variational parameters of local variables were optimized using a coordinate ascent algorithm. One of the most difficult problems is the intractable computation of expectations in equation 6. Depending on the framework of the mixture models specifically considered for each omics dataset, several special expectations cannot be obtained directly from analytically tractable solutions. For example, the DMM showed that the expectations of the logarithms of the multinomial beta function could not be calculated analytically. To overcome these problems, mathematical expansions such as the Taylor expansion and the delta method were adopted in this study. Mathematical explanations of these expansions and variational objective functions are provided in the Supplementary Material.

In particular, the dimensionality of integrated microbiome multi-omics data sets can rapidly increase if the number of different omics datasets increases. For example, the average number of species in the microbiome data is approximately tens of thousands, and metabolomics datasets include thousands of metabolite profile features. Thus, the variational parameters of the global variables were optimized using stochastic algorithms. The stochastic inference is much more computationally efficient because it updates variational factors by sampling the data points in each iteration and uses the natural gradient method, which can achieve faster convergence than standard gradients [21].

Figure 1 shows workflow schematics of the I-SVVS approach. The input to I-SVVS consists of matrices of metabolite profiles, which are continuous variables, and microbiome species, which are count variables (Fig. 1a). I-SVVS uses the hierarchical Dirichlet mixture process to integrate metabolite profile data and microbiome species (or taxonomic unit) data from multi-omics experiments as a composition of biological and technical sources of variation. I-SVVS can identify important features in the metabolite profiles and microbiome species data for each cluster. Because the distributions of latent variables are intractable when computing their posteriors, the I-SVVS approach optimizes the parameters of both of components simultaneously using the SVI approach [31]. To overcome the extra-dimensional problems of integrating metabolite profiles and microbiome species databases, highly efficient techniques for stochastic optimization were adopted for the computational process. The output of the I-SVVS approach consists of two main components (Fig. 1b). The first component clustered all samples by integrating the metabolite profile and microbiome species data. The second component provides an important core set of microbiome species (or taxonomic units) and metabolite profiles that significantly contribute to the clustering process. Due to the integration of metabolite profiles and microbiome data, shared information influenced the variable selection of the two databases. Following variational inference, the I-SVVS results were used for subsequent analysis. For example, integrated clustering results can be used for low-dimensional visualizations such as principal-coordinate analysis (PCoA) and nonmetric multidimensional scaling (NMDS) [32]. The results of the variable selection of microbiome species and metabolites were used for phylogenetic and metabolic network analyses, respectively.

**Fig 1.**
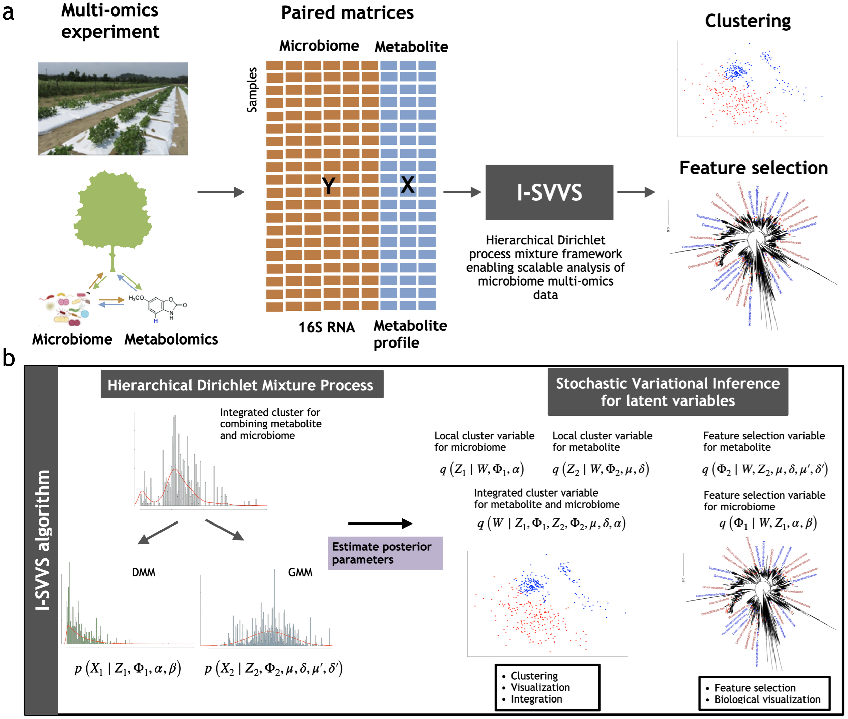
Schematic of a microbiome multi-omics data analysis pipeline with I-SVVS. **a**, An example of a microbiome multi-omics experiment on a plant simultaneously measures metabolite profile and microbiome species in each sample, producing paired matrices for metabolite and microbiome. These matrices include a matrix of count values for microbiome species data and another matrix of continuous values for metabolite profile data. They are the input to the I-SVVS, which integrates all databases to cluster samples and selects the important features of each database. **b**, Schematic of the I-SVVS approach. Firstly, the infinite Dirichlet multinomial mixture (DMM) model with variable selection is proposed for microbiome data, *Z*_1_ is an allocation variable, Φ_1_ is a feature selection variable, *α* is parameter of Dirichlet-multinomial distribution for the selected group of microbiome species and *β* is parameter of Dirichlet-multinomial distribution for the rejected group of microbiome species. The infinite Gaussian mixture model (GMM) with variable selection is proposed for the metabolite profile data, *Z*_2_ is an allocation variable, Φ_2_ is a feature selection variable, *µ, δ* are the parameters of normal distribution captured the selected group of metabolite profiles and *µ*^*′*^, *δ*^*′*^ are parameters of normal distribution captured the rejected group of metabolite profiles. Then the framework of the hierarchical Dirichlet mixture process integrates the information of GMM and DMM approaches that allow the shared information between metabolite profile and microbiome data for the clustering process. W denotes the integrated (global) allocation variable. Next, the stochastic variable inference is proposed to estimate the posterior distributions of the latent variables for clustering and variable selection (Methods).

## Database description

### Study inclusion and data acquisition

Dataset A represents the environmental microbiome data from the soybean field experiments, which included 186 drought irrigation samples, 191 control samples, 16,943 microbiome species (or taxonomic units), and 265 metabolome features. The experimental explanations for Dataset A are provided in the Supplementary Material.

We also employed case-control studies from two published gut microbiome and metabolome datasets in mice and humans. Dataset B represents the obstructive sleep apnea (OSA) dataset for mice, which includes 102 samples for intermittent hypoxia and hypercapnia (IHH), 102 samples for control, 4,690 taxonomic units and 1,710 metabolome features [29]. Dataset C represents a study on C. difficile infection (CDI) in humans. The 338 samples included 3,347 taxonomic units and 103 metabolome features [30].

### Open-source software

The software is implemented in Python and used standard libraries, such as NumPy and SciPy, for mathematical computations. The software inputs microbiome count data and metabolites data in a CSV file and outputs the inferred clusters and a core set of selected taxonomic units and metabolism features. The main options in the software tool are the maximum number of global and local clusters, which pose limitations in estimating the number of clusters, and the number of taxonomic units and metabolism features that users want to select. I-SVVS uses the iterative optimization algorithms to estimate the parameters; thus, a convergence criterion is used to implement a stopping rule [21]. The I-SVVS algorithm stops when the change in the ELBO computations is less than 1e-3 (Supplementary Material). We use the convergence criterion fixed across all datasets in this study. The number of iterations should be modified for datasets notably smaller or larger in scale than those considered in this study. This is a tunable option in the software. The software is available at https://github.com/tungtokyo1108/I-SVVS.

Similar to our previous research, we set the cluster count truncation levels for global and local clusters to 10 and hyperparameters of the stick-breaking representations to 0.1 during initialization [21]. A comprehensive explanation of the initial values for the hyperparameters of all priors can be found in the supplementary materials.

To tackle the selection of taxonomic units and metabolism features using the I-SVVS, we compute the averages of Φ_1_ and Φ_2_ across samples after estimating their values, respectively. Subsequently, we arrange taxonomic units and metabolism features in descending order based on these averaged Φ_1_ and Φ_2_ values. Our software package generates tables that present these ranked values, enabling users to choose a core set of taxonomic units and metabolism features from the top values in these tables.

## Results

### I-SVVS accurately clusters large datasets by integrating multiple data types

The application of I-SVVS to multi-omics microbiome data shows that the global cluster assignment variable helps share information between microbiome species and metabolite profiles to significantly improve the performance of the clustering process. Moreover, the selected features of the two datasets provide excellent interpretations of the obtained clusters for biological exploration.

To illustrate this, we applied I-SVVS to three multiomics microbiome datasets from humans, mice, and plants, spanning thousands to tens of thousands of features from microbiome species and metabolite profiles. We compared the performances of I-SVVS with the integration of microbiome and metabolomics data with the DMM approach [20, 21] with only microbiome species data, as well as with iClusterPlus [17] which is a general-purpose clustering method commonly applied to multi-omics data. The current version of iClusterPlus has been developed in a framework that allows the integration of categorical, count, and continuous data, and can thus analyze the integration of metabolite profile data and microbiome species data. Moreover, Clusternomics, a probabilistic clustering method [16], was employed to compare the performance of I-SVVS. The agreement between the clustering obtained with the three approaches and the ground-truth clustering was measured using the Adjusted Rand Index (ARI) used in our previous study [21]. We followed the Deviance Information Criterion (DIC) for the selection of Bayesian models and default the values of the Clusternomics 0.1.1 packages in R to determine the number of clusters [16]. iClusterPlus uses a deviance ratio metric, which can be interpreted as the percentage of the total variation, to select the number of clusters. We followed the default values of the *tune*.*iClusterPlus* function in the iClusterPlus 1.32.0 package in R and selected a Gaussian distribution for the metabolic data and a Poisson distribution for the microbiome data [17].

Table 1 presents the computational time required for the calculations of I-SVVS, iClusterPlus, and Clusternomics. We found that the I-SVVS was able to significantly reduce the running times for Datasets A, B, and C. The computational time was found to increase considerably when the number of taxonomic units became very large, such as in dataset A. Moreover, although the number of features of metabolome profile data in dataset B (1,710 features) was substantially larger than that in dataset C (103 features), there was a small difference in the number of taxonomic units between the two datasets. Table 1 shows that the computational time for Dataset C is trivially faster than that for Dataset B. Therefore, the high dimensionality of microbiome data is the most important factor influencing the computational burden of multi-omics analysis. We observed that the scalability of the I-SVVS approach can significantly reduce the computational time required to handle very high-dimensional datasets; for instance, processing a complete set of approximately 17,000 microbiome species and 260 metabolic features in approximately 2 h.

**Table 1.** Running time of the three approaches on the real data sets.

| Dataset | Clusternomics | iClusterPlus | I-SVVS |
| --- | --- | --- | --- |
| A | 2.35 d | 1.12 d | 2.18 h |
| B | 13.12 h | 6.26 h | 18.42 min |
| C | 10.54 h | 4.78 h | 15.36 min |

Note: All algorithms were run on a personal computer (Intel® Xeon® Gold 6230 Processor 2.10 GHz × 2, 40 cores, 2 threads per core, 128 Gb RAM) under Ubuntu 20.04.1 LTS.

Next, we calculated the Adjusted Rand Index (ARI) to evaluate the performance of the three approaches with the three datasets. Table 2 shows that the I-SVVS approach achieved the best performance for datasets A, B, and C (ARI was 0.89, 0.78, and 0.73 for datasets A, B, and C respectively). The ARI value of I-SVVS was highest in Dataset A, which had the largest number of taxonomic units. Because I-SVVS integrates an infinite Dirichlet multinomial mixture, which is a specific model for analyzing microbiome OTU data [21], this approach achieved better performance than other approaches that were not developed to analyze microbiome data in the process. Table 2 shows that the ARI value of iClusterPlus was highest for Dataset A (ARI = 0.81). In addition, I-SVVS, iClusterPlus, and Clusternomics exhibited poor performance on dataset C (ARI = 0.73, 0.6, and 0.48, respectively). The main reason for this could be that the number of features in the metabolome profile of dataset C was significantly smaller than that of the others. Figure 2 shows the confusion matrix plots for Dataset A, calculated using I-SVVS, iClusterPlus, and Clusternomics. The drought and control groups were clustered using I-SVVS with accuracies of 90% and 88%, respectively. Figures S1a-c show the estimated mixing coefficients of the clusters in datasets A, B, and C after convergence. In some clusters, the estimated mixing coefficients were close to zero after convergence. Therefore, a highly likely number of clusters can be obtained for the mixtures. Figure S1a shows the strongest support for the two clusters in dataset A because the mixing coefficients of clusters 3 and 4 have larger values, while Figure S1b shows the highest probability of the two clusters in dataset B because the mixing coefficients of clusters 1 and 4 have large values; Figure S1c shows the highest probability of 3 clusters in dataset C.

**Table 2.** ARI scores of the three approaches on the real data sets.

| Dataset | Clusternomics | iClusterPlus | I-SVVS |
| --- | --- | --- | --- |
| A | 0.722 | 0.815 | 0.891 |
| B | 0.553 | 0.697 | 0.781 |
| C | 0.482 | 0.602 | 0.732 |

**Fig 2.**
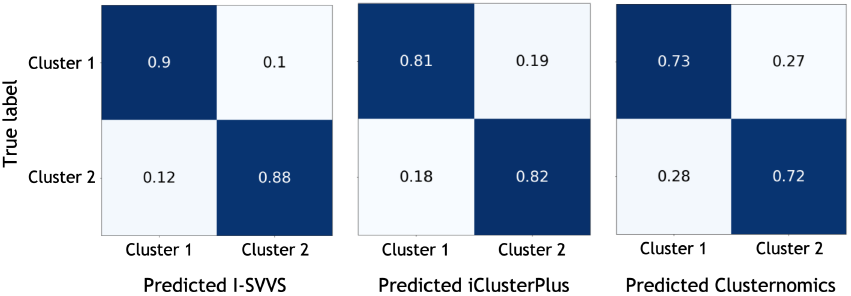
Confusion matrix plots of dataset A with labels indicating predicted class using the three approaches and true group. Cluster 1 denotes the drought and cluster 2 denotes the control. (Left) I-SVVS, (Middle) iClusterPlus, (Right) Clusternomics

### I-SVVS identifies a core set of features for microbiome species and metabolite profiles

Several previous studies have investigated the important roles of metabolism-microbiome associations in mice [33, 34, 35], and plants [36]. To address this trend, the I-SVVS approach was used to identify a core set of microbial species and metabolic features that showed significant differences among the clusters obtained in the analysis. Figures 3a-b show the histograms of the averages of microbiome indicator variable Φ_1*ij*_ and metabolism indicator variable Φ_2*ij*_*′* over *i*^*th*^ sample in dataset A. The distribution depicted in Figure 3a showcases prominent peaks centered around 0.6, devoid of any conspicuous outliers. Microbiome species with Φ_1*ij*_ values of 0.8 demonstrate significant contributions to the classification process. The I-SVVS method assigns substantial weights to microbiome species exhibiting remarkable signals. The Φ_1*ij*_ values nearing 0.6 may suggest a lack of robust signaling in microbiome species, indicating a mild or weak influence on classification. Consequently, such species undergo appropriate down-weighting. The current model adeptly diminishes the impact of microbiome species with weak signals. Moreover, microbiome species with Φ_1*ij*_ values below 0.3 are prevalent among groups with minimal contributions to classification, underscoring the effectiveness of I-SVVS in discerning and prioritizing pivotal microbiome species. Similarly, the distribution portrayed in Figure 3b exhibit prominent peaks centered around 0.55, without any glaring outliers. Metabolome profile features characterized by Φ_2*ij*_*′* values of 0.7 contribute significantly to the classification process. The I-SVVS methodology assigns substantial weights to metabolome profile features displaying remarkable signals. The Φ_2*ij*_*′* values nearing 0.55 may suggest a lack of robust signaling in metabolome profile features, indicating a mild or weak influence on classification. As a result, such features undergo appropriate down-weighting. The current model skillfully reduces the impact of metabolome profile features with weak signals. Additionally, metabolome profile features with Φ_2*ij*_*′* values below 0.3 are prevalent among groups with minimal contributions to classification, underscoring the effectiveness of I-SVVS in discerning and prioritizing crucial metabolome profile features. Figure 4 shows that the top 100 selected microbial species in dataset A were mapped on the 16S phylogenetic tree. The identification of group-microbiome and group-metabolism associations is based on Φ_1*ij*_ and Φ_2*ij*_*′* that *i*^*th*^ sample is assigned to a specific cluster (drought and control conditions) in dataset A. As expected, our results are consistent with those of our previous study that analyzed soybean rhizosphere microbiome data. For example, I-SVVS selected important microbiome families that were significantly associated with plant growth promotion under control conditions, such as the *Chitinophagaceae* family within the order *Chitinophagales* from the phyla *Bacteroidetes, Nitrosomonadaceae* and *Chromobacteriaceae* families within the order *Gammaproteobacteria* from the phylum *Proteobacteria*. These are the dominant bacterial phyla in the soybean rhizosphere [37, 38]. Several studies showed [39, 40] the *Nitrosomonadaceae* family oxidize ammonia to nitrite using the enzyme ammonia monooxygenase (AMO), which catalyzes the first step of ammonia oxidation. This family is a key group of nitrifying bacteria that plays a vital role in the conversion of nitrogen compounds in natural ecosystems and agricultural systems [41]. *Chitinophagaceae* family produces enzymes called chitinases that are responsible for the degradation of chitin [42, 43]. The enzymatic breakdown of chitin into smaller components can be further metabolized into a source of energy, carbon, and nitrogen. This capability is important for protecting plants against fungal infections [43]. Moreover, our study showed that the *Microbacteriaceae* family within the order *Micrococcales* and the *Micromonosporaceae* family within the order *Micromonosporales* from the phylum *Actinobacteriota, Beijerinckiaceae*, and *Hyphomicrobiaceae* families within the order *Rhizobiales* from the class *Alphaproteobacteria* were more abundant in the drought treatments. Several studies have shown that the *Microbacteriaceae* and *Micromonosporaceae* families can improve the growth of plant hosts through nitrogen fixation, which converts dinitrogen (*N*_2_) into ammonia [44, 45, 46]. Therefore, the host plant can utilize these natural sources of nitrogen and reduce its dependence on external sources, such as fertilizers.

**Fig 3.**
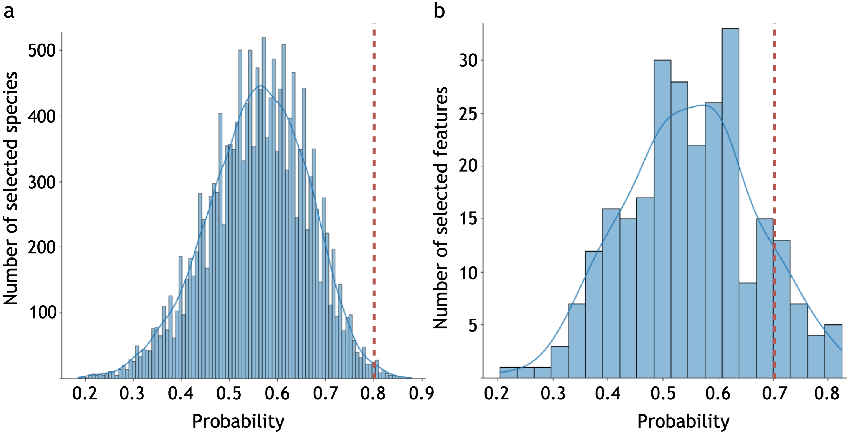
Histogram of the average of Φ_1_ and Φ_2_ in dataset A. The dashed lines are bound to select microbiome species and metabolome profile features. a. microbiome data; b. metabolome data

**Fig 4.**
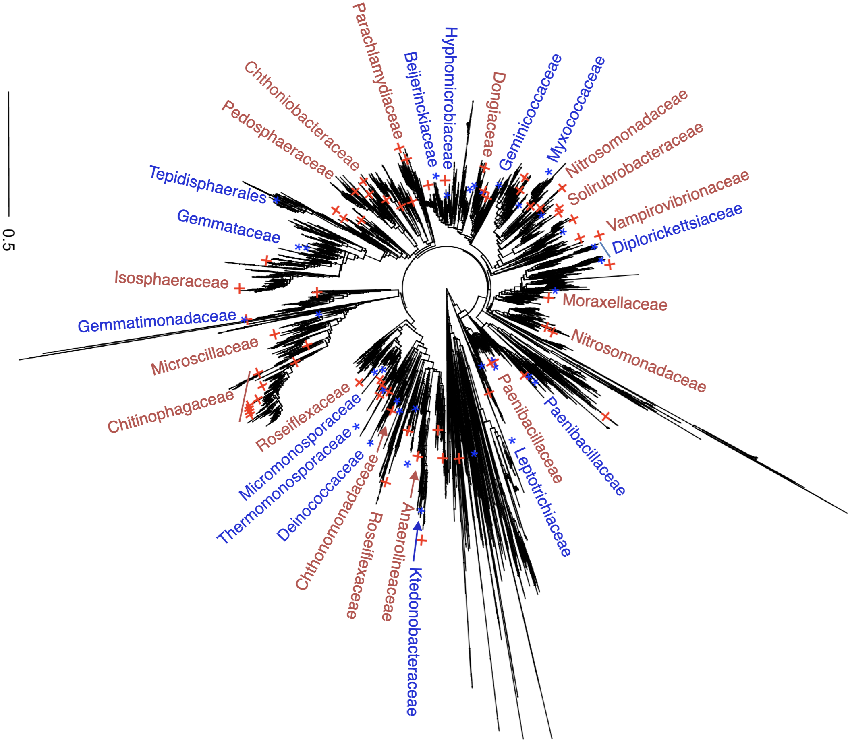
Microbial species selected using the integrative stochastic variational variable selection (I-SVVS) approach and mapped on the phylogenetic tree for dataset A. Red-colored plus symbols denote the control and blue-colored starts denote drought

To assess large-scale metabolite production and consumption patterns, we hierarchically clustered the top individual metabolism and microbiome that were selected by the I-SVVS approach into two groups of dataset A (figure 5). Our analysis revealed significant correlations between microbiome species (Φ_1_) and metabolic traits (Φ_2_) across subjects in the control and drought groups such as 1-aminocyclopropane-1-carboxylate (ACC), L-proline, L-tyrosine, L-aspartic acid, and L-glutamic acid. Previous studies have shown that 1-aminocyclopropane-1-carboxylate (ACC), an important intermediate in the ethylene synthesis, can change the soil microbiome to enhance plant tolerance to salinity stress. A group of beneficial bacteria can degrade ACC to ammonia and *α*-ketobutyrate using ACC deaminase, thereby decreasing the level of ethylene and enhancing plant growth [47, 48]. Numerous investigations have elucidated the profound influence of glutamic acid on restructuring the plant microbial community. This influence manifests in the augmentation of population sizes not only within *Streptomyces* but also among *Bacillaceae* and *Burkholderiaceae*, thereby mitigating disease incidence as indicated by previous studies [49, 50]. Intriguingly, glutamic acid’s lack of activation of host plant resistance mechanisms implies its potential to unveil pivotal insights into the evolutionary and functional dynamics between the plant and its microbiota. Particularly noteworthy is the metabolic utilization of glutamic acid by *Streptomyces* as the sole source of carbon and nitrogen, offering a plausible explanation for its discernible impact on the interconnected plant-associated communities [51]. One strategy employed by beneficial bacteria in the rhizosphere to dampen plant immunity involves the biosynthesis of gluconic acid. Notably, bacterial strains like *Pseudomonas capeferrum* and *Pseudomonas aeruginosa* produce gluconic acid, contributing to a reduction in rhizosphere pH. This pH alteration, in turn, acts as a mechanism to suppress plant immunity, highlighting the intricate ways in which beneficial bacteria interact with the plant environment. [52].

**Fig 5.**
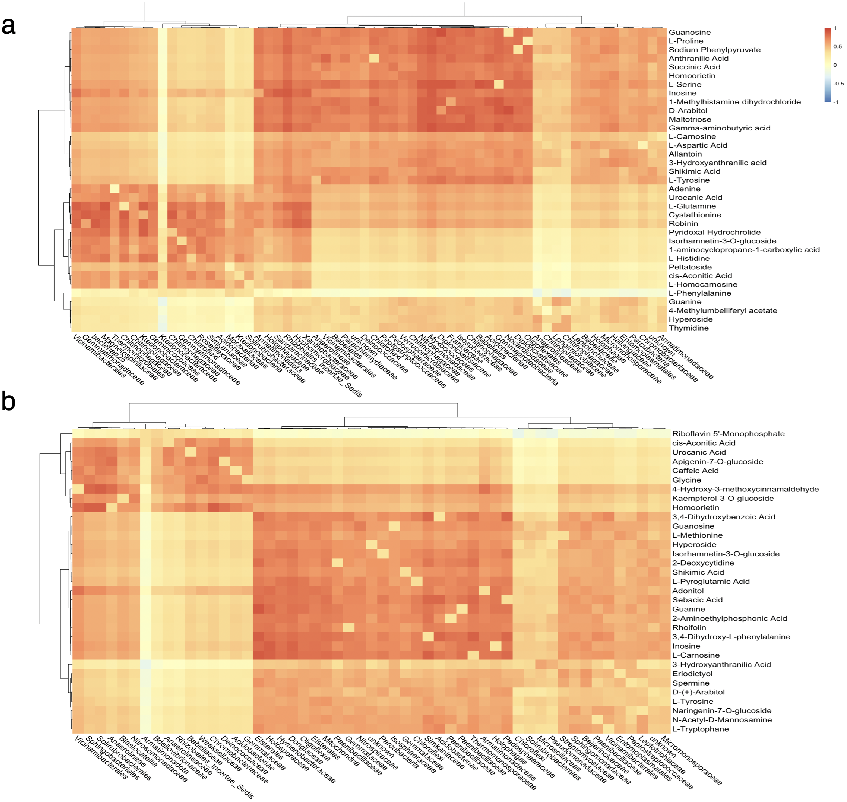
Heat map of microbiome species (Φ_1_) and metabolic traits (Φ_2_) that were selected by the I-SVVS approach in two groups for dataset A. Individual metabolites and microbiomes were hierarchically clustered (Ward’s method) using Euclidean distance. a. control group; b. drought group

In the mouse fecal samples of obstructive sleep apnea (OSA) in dataset B, Figure S2a-b show the histograms of the averages of the microbiome indicator variable Φ_1*ij*_ and the metabolism indicator variable Φ_2*ij*_*′* over *i*^*th*^ sample in dataset B. The distribution illustrated in Figure S2a showcases prominent peaks centered around 0.5, without any conspicuous outliers. Microbiome species with Φ_1*ij*_ values of 0.81 exhibit significant contributions to the classification process. The I-SVVS method allocates substantial weights to microbiome species displaying remarkable signals. The Φ_1*ij*_ values nearing 0.5 may indicate a lack of robust signaling in microbiome species, suggesting a mild or weak influence on classification. Consequently, such species undergo appropriate down-weighting. The current model adeptly mitigates the impact of microbiome species with weak signals. Furthermore, microbiome species with Φ_1*ij*_ values below 0.3 are prevalent among groups with minimal contributions to classification, underscoring the effectiveness of I-SVVS in discerning and prioritizing pivotal microbiome species. In the same way, the distribution portrayed in Figure S2b exhibit prominent peaks centered around 0.5, lacking any glaring outliers. Metabolome profile features characterized by Φ_2*ij*_*′* values of 0.8 contribute significantly to the classification process. The I-SVVS methodology assigns substantial weights to metabolome profile features displaying remarkable signals. The Φ_2*ij*_*′* values nearing 0.5 may suggest a lack of robust signaling in metabolome profile features, indicating a mild or weak influence on classification. As a result, such features undergo appropriate down-weighting. The current model diminishes the impact of metabolome profile features with weak signals. Additionally, metabolome profile features with Φ_2*ij*_*′* values below 0.3 are prevalent among groups with minimal contributions to classification, underscoring the effectiveness of I-SVVS in discerning and prioritizing crucial metabolome profile features. Figure S3 shows that the top 100 selected microbial species in dataset B were mapped on the 16S phylogenetic tree. The identification of group-microbiome and group-metabolism associations was based on Φ_1*ij*_ and Φ_2*ij*_*′*, where the *i*^*th*^ sample was assigned to a specific cluster (intermittent hypoxia and hypercapnia (IHH) cases and air controls) in dataset B. I-SVVS selected important microbiome families that were significantly associated with IHH exposure, such as *Lachnospiraceae* and *Ruminococcaceae* families in the phylum *Firmicutes*. These results were also reported in previous study [29]. Previous studies on sleep fragmentation showed that the growth of highly fermentative members of the *Lachnospiraceae* and *Ruminococcaceae* families lead to visceral white adipose tissue inflammation and alterations in insulin sensitivity [53, 54]. Moreover, I-SVVS identified several important metabolomic features, such as chenodeoxycholic acid and cholic acid, which were significantly associated with IHH exposure. These are primary bile acids that play important roles in facilitating the digestion and absorption of cholesterol and triglycerides. Several studies have reported alterations in response to intermittent hypoxia [55, 56].

## Discussion

Integrative stochastic variational variable selection (I-SVVS) is a Bayesian nonparametric method that uses an integrated multi-omics dataset of the microbiome to rapidly cluster samples and select important features from multi-omics data. We applied I-SVVS to extreme-dimensional microbiome multi-omics profiles collected in our own soybean experiments and published datasets of mice and humans. I-SVVS is also unique in its ability to integrate the microbiome dataset (such as 16S rRNA) with metabolome, which is not possible with current approaches such as iClusterPlus and Clusternomics.

First, in the soybean dataset (Dataset A), we demonstrated that I-SVVS can achieve accurate clustering based on the integration of the metabolite profile and microbiome data. Owing to the hierarchical Dirichlet mixture models, I-SVVS can capture not only information shared by microbiome and metabolome datasets but also those emerging from the complementarity of these datasets. Our results also demonstrate that I-SVVS can leverage information from multiple omics layers to accurately cluster samples from sparse profiling datasets and avoid the problem of instability of inferred clusters in previous probabilistic algorithms. Most notably, I-SVVS identified an important core set of representative features that vary per sample rather than per cluster from a large number of multi-omics biological features, that is, the microbiome and metabolome. Identification cannot be conducted using the previous Bayesian methods such as BCC and Clusternomics. Our previous method (SVVS) [21] identified the important features (taxonomic units) using only microbiome data. Therefore, I-SVVS-supported selected features play an important role in significantly improving the performance of clustering analysis and interpreting shared information and interactions between different types of omics data. For example, I-SVVS can be used to investigate the relationship between metabolites and the microbiome community structure and function, which plays a crucial role in studies on human health and disease [57, 58]. Moreover, to overcome the computational burden of high-dimensional microbiome multi-omics data, I-SVVS uses stochastic variational inference to estimate a model parameters. We also applied I-SVVS to a joint analysis of multi-omics microbiome datasets from mice and humans. In these datasets, I-SVVS achieved good performance in identifying the key features that highlighted the impact of disease on host-commensal organism co-metabolism in human and animal guts. Therefore, the flexibility and scalability of I-SVVS make it easily applicable to multiple datasets with larger dimensionality and enable extensions that incorporate additional omic technologies.

Although we proposed several solutions to overcome important challenges in microbiome multi-omics analysis, I-SVVS is not free of limitations. The model focuses on optimizing the contributions of microbiome data, which could significantly improve its performance in the joint analysis of multi-omics datasets. Although the DMM approach is the best mixture model for analyzing microbiome count data, it cannot efficiently model count data of different omics data. Future extensions of I-SVVS may address problems that develop and integrate specific Bayesian mixture models of different omics data, such as metatranscriptome RNA sequencing (MT), and shotgun mass spectrometry-based metaproteomics (MP) [59] in its framework. In addition, variations in the structure of omics data, such as an imbalance in the number of features of each omics dataset, affect the stability and optimal performance of I-SVVS. Future research should also consider this point. Finally, although I-SVVS successfully identified a small number of vital features in different omics datasets, it was difficult to infer the interactions among the selected features. Therefore, there is room for future extensions that are more efficient at enforcing the important relationships across omics than the use of correlations.

## Conclusion

In conclusion, the proposed integrative stochastic variational variable selection approach has the potential to significantly enhance the effectiveness of the Bayesian mixture model for joint analysis of high-dimensional multi-omics microbiome data. The selected minimal core set of microbial species and metabolites simplifies the identification of key features that have the greatest impact on the distinctions among samples. This study will make a significant contribution to and inspire continued endeavors aimed at enhancing the efficiency of Bayesian statistical models for the rapid identification of crucial features within multi-omics microbiome data across various domains of research.

## Author contributions statement

T.D. and H.I. designed the study. Y.T., Y.Y., H.T. and H.I. designed and conducted the field experiment in Tottori. K.K., E.U., S.K., T.S. and Y.I. performed the microbiome analysis from tissue sampling, library preparation, sequencing, and primary informatics for taxonomic assignment and diversity statistics. T.D. developed the method and analyze the data. T.D. and H.I. interpreted the data and wrote the manuscript. The authors read and approved the final manuscript.

## Acknowledgments

We are grateful to the technical staff of the Arid Land Research Center, Tottori University, and Izumi Higashida for managing of the field experiments on soybean. We would like to thank all the members of the JST-CREST Program including Mikio Nakazono, Hirokazu Takahashi, Toru Fujiwara, Yoshihiro Ohmori, Hideki Takanashi, Akito Kaga, Mai Tsuda and Yuji Sawada for conducting the field experiments.

This work was supported by JSPS KAKENHI (Grant Number JP21J21850), the JST-CRESET Program (Grant Number JPMJCR1602), the JST-Mirai Program (Grant Number JPMJMI120C7), and the JST ALCA-Next Program (Grant Number JPMJAN23D1), Japan.

