## Supplementary for "I-SVVS: Integrative stochastic variational variable selection to explore joint patterns of multi-omics microbiome data"

### 1 Supplementary Figures

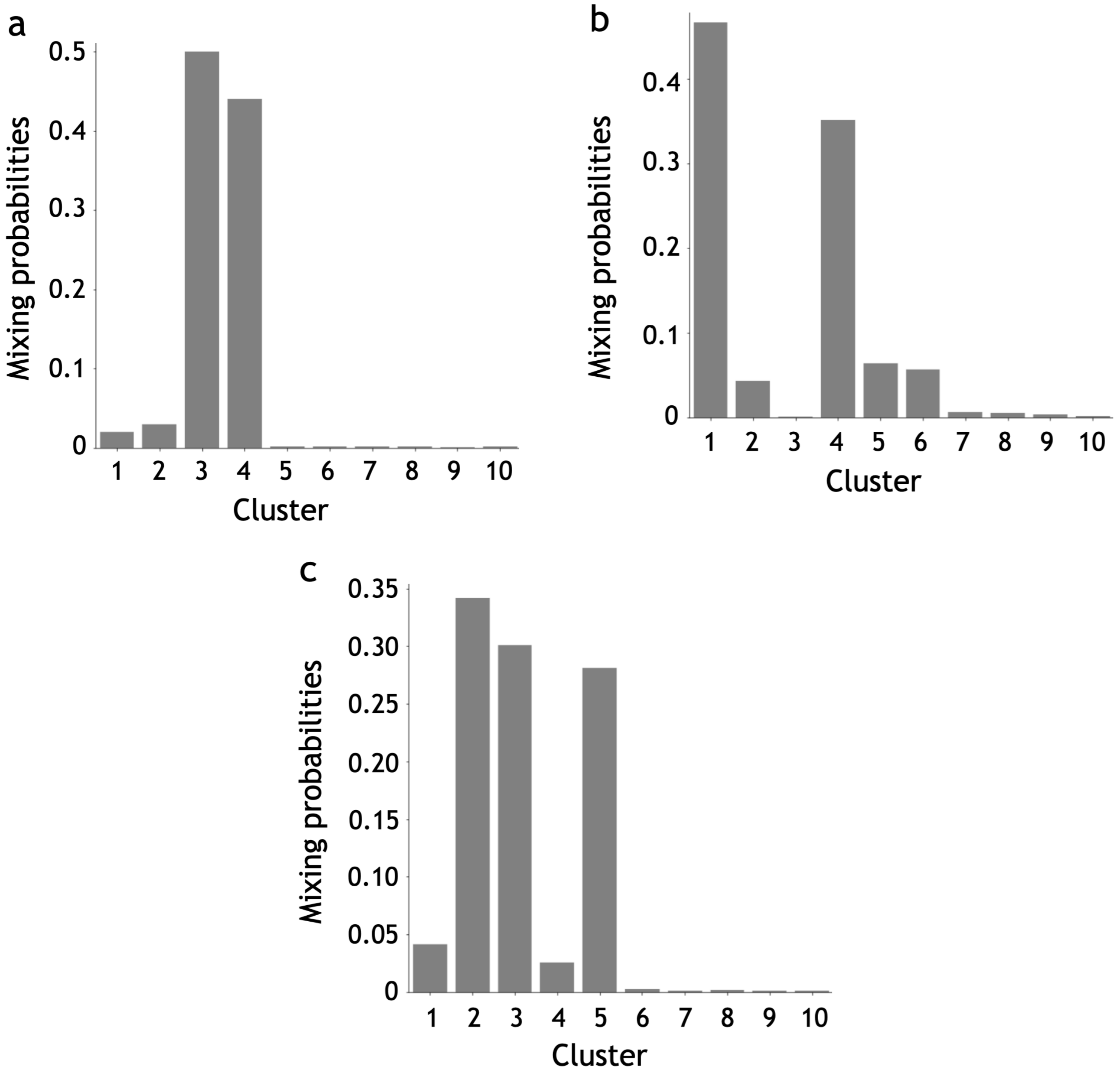


**Supplementary Figure S1**: The estimated values of the mixing coefficients. a. dataset A; b. dataset B; c. dataset C

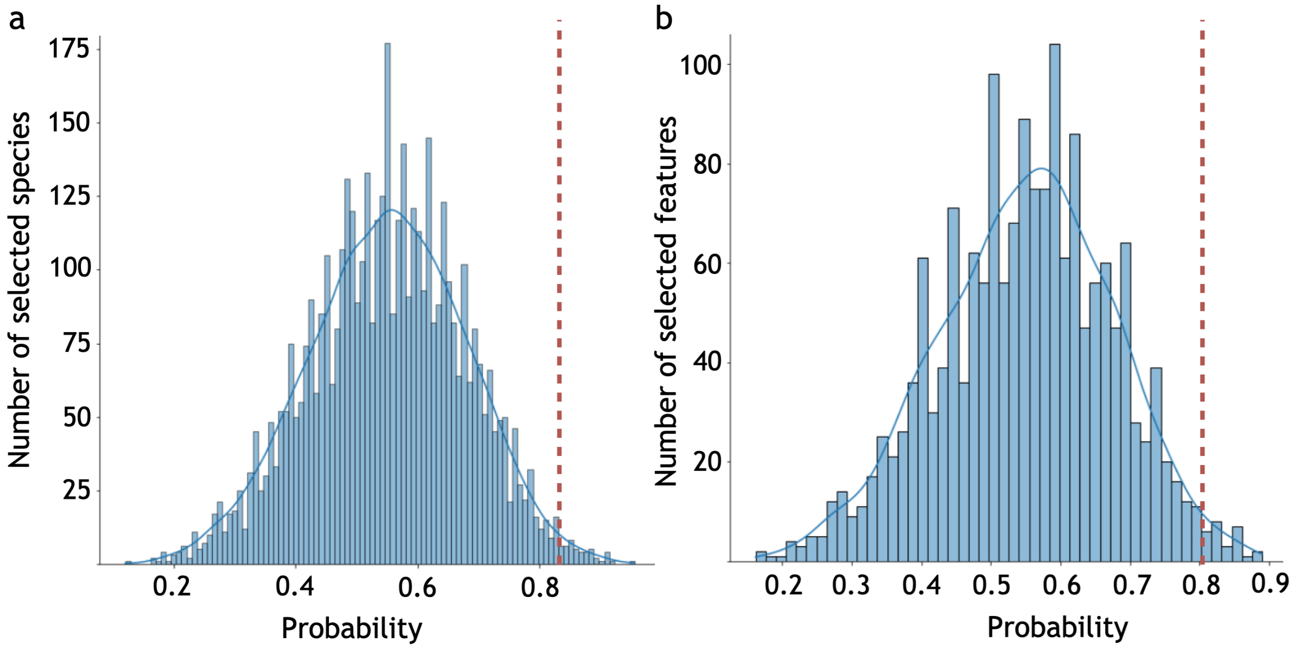


**Supplementary Figure S2**: Histogram of the average of $\Phi_1$ and $\Phi_2$ in dataset B. The dashed lines are bound to select microbiome species and metabolome profile features. a. microbiome data; b. metabolome data.

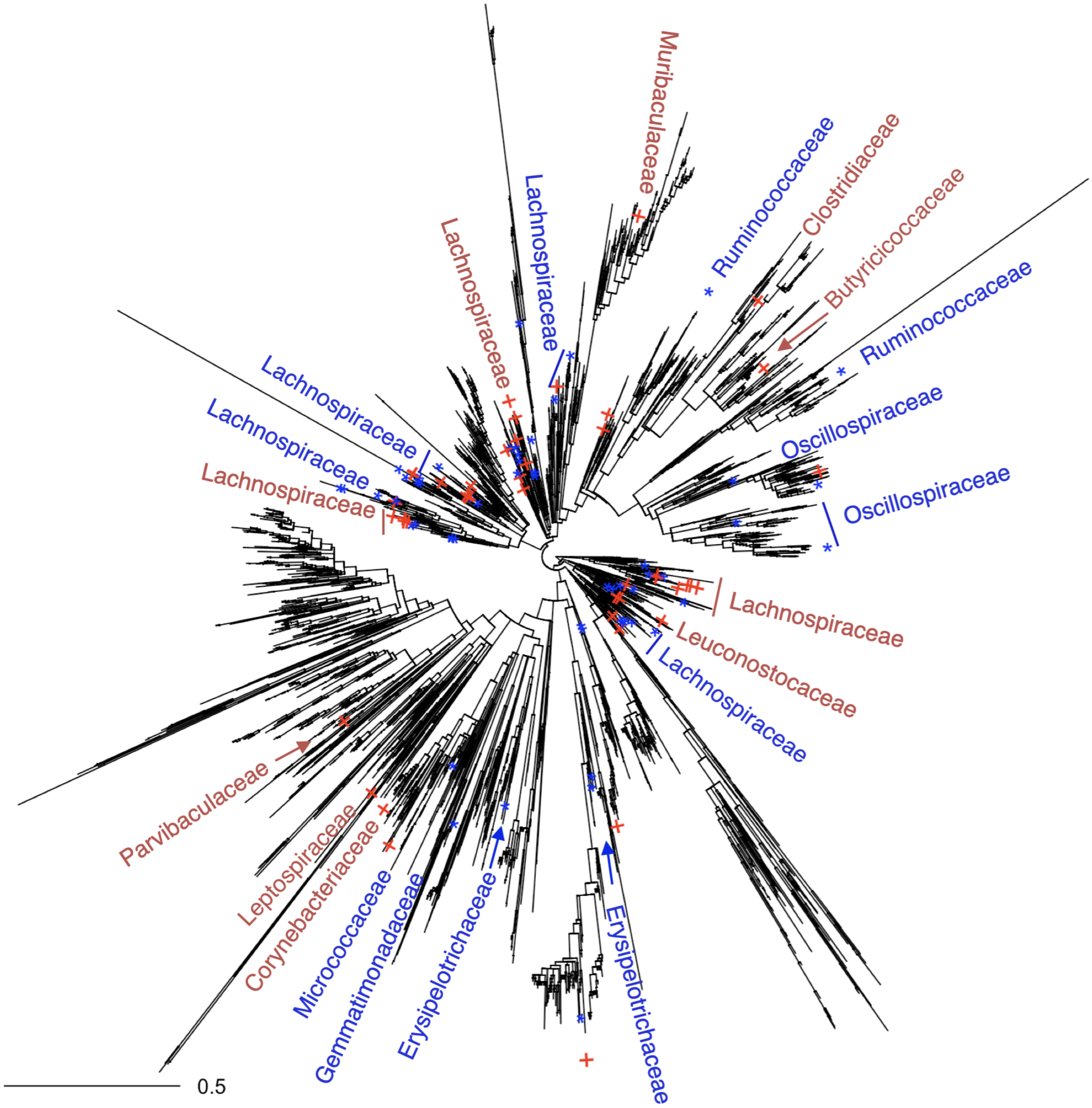


**Supplementary Figure S3**: Microbial species selected using the integrative stochastic variational variable selection (I-SVVS) approach and mapped on the phylogenetic tree for dataset B. Blue-colored starts symbols denote intermittent hypoxia and hypercapnia (IHH) cases and Red-colored plus denote air controls.

#### 2 Mean-Field Variational Inference for a integration of Gaussian mixture model (GMM) and Dirichlet multinomial mixtures (DMM) Model

We expand specifically the variational lower bound equation as follows:

$$
\begin{aligned}
\mathcal{L} = & \sum_{k=1}^{K_{max}} \mathrm{E}_q \left[\log\left(p\left(\Psi_k^{'}|1,\kappa\right)\right)\right] - \sum_{k=1}^{K_{\max}} \mathrm{E}_q \left[\log\left(q\left(\Psi_k^{'}|\vartheta_k,\vartheta_k^{'}\right)\right)\right] \\
& + \sum_{i=1}^{N}\sum_{k=1}^{K_{max}} \mathrm{E}_q \left[\log p(W_i^k|\Psi_1^{'},\Psi_2^{'},...,\Psi_{K_{max}}^{'})\right] - \sum_{i=1}^{N}\sum_{k=1}^{K_{\max}} \mathrm{E}_q \left[\log\left(q\left(W_i^k|g_i^k\right)\right)\right] \\
& + \sum_{t=1}^{T_{max}} \mathrm{E}_q \left[\log\left(p\left(\Pi_{1t}^{'}|1,\kappa^*\right)\right)\right] - \sum_{t=1}^{T_{\max}} \mathrm{E}_q \left[\log\left(q\left(\Pi_{1t}^{'}|\vartheta_t^*,\vartheta_t^{*'}\right)\right)\right] \\
& + \sum_{k=1}^{K_{max}}\sum_{t=1}^{T_{max}} \mathrm{E}_q \left[\log p(Z_{1t}^k|\Pi_{11}^{'},\Pi_{12}^{'},...,\Pi_{1T_{max}}^{'})\right] - \sum_{k=1}^{K_{\max}}\sum_{t=1}^{T_{max}} \mathrm{E}_q \left[\log\left(q\left(Z_{1t}^k|l_{1t}^k\right)\right)\right] \\
& + \sum_{i=1}^{N}\sum_{j=1}^{D} \mathrm{E}_q \left[\log p(\Phi_{1ij}|\epsilon_{j_1},\epsilon_{j_2})\right] - \sum_{i=1}^{N}\sum_{j=1}^{D} \mathrm{E}_q \left[\log\left(q\left(\Phi_{1ij}|f_{ij}\right)\right)\right] \\
& + \sum_{j}^{D} \mathrm{E}_q \left[\log\left(p\left(\epsilon_j|\xi\right)\right)\right] - \sum_{j}^{D} \mathrm{E}_q \left[\log\left(q\left(\epsilon_j|\xi^*\right)\right)\right] \\
& + \sum_{j=1}^{D}\sum_{t=1}^{T_{max}} \mathrm{E}_q \left[\log\left(p\left(\alpha_{jt}|\lambda_{jt}\right)\right)\right] - \sum_{j=1}^{D}\sum_{t=1}^{T_{max}} \mathrm{E}_q \left[\log\left(q\left(\alpha_{jt}|\lambda_{jt}^*\right)\right)\right] \\
& + \sum_{j=1}^{D} \mathrm{E}_q \left[\log\left(p\left(\beta_j|\iota_j\right)\right)\right] - \sum_{j=1}^{D} \mathrm{E}_q \left[\log\left(q\left(\beta_j|\iota_j^*\right)\right)\right] \\
& + \sum_{t^{'}=1}^{T_{max}^{'}} \mathrm{E}_q \left[\log\left(p\left(\Pi_{2t^{'}}^{'}|1,\kappa^+\right)\right)\right] - \sum_{t^{'}=1}^{T_{\max}^{'}} \mathrm{E}_q \left[\log\left(q\left(\Pi_{2t^{'}}^{'}|\vartheta_t^+,\vartheta_t^{+'}\right)\right)\right] \\
& + \sum_{k=1}^{K_{max}}\sum_{t^{'}=1}^{T_{max}^{'}} \mathrm{E}_q \left[\log p(Z_{2t^{'}}^k|\Pi_{21}^{'},\Pi_{22}^{'},...,\Pi_{2T_{max}^{'}}^{'})\right] - \sum_{k=1}^{K_{\max}}\sum_{t^{'}=1}^{T_{max}^{'}} \mathrm{E}_q \left[\log\left(q\left(Z_{2t^{'}}^k|l_{2t^{'}}^k\right)\right)\right] \\
& + \sum_{i=1}^{N}\sum_{j^{'}=1}^{D^{'}} \mathrm{E}_q \left[\log p(\Phi_{2ij^{'}}|\epsilon_{j_1^{'}}^{'},\epsilon_{j_2^{'}}^{'})\right] - \sum_{i=1}^{N}\sum_{j^{'}=1}^{D^{'}} \mathrm{E}_q \left[\log\left(q\left(\Phi_{2ij^{'}}|f_{ij^{'}}^{'}\right)\right)\right] \\
& + \sum_{j^{'}}^{D^{'}} \mathrm{E}_q \left[\log\left(p\left(\epsilon_{j^{'}}^{'}|\xi^{'}\right)\right)\right] - \sum_{j^{'}}^{D} \mathrm{E}_q \left[\log\left(q\left(\epsilon_{j^{'}}^{'}|\xi^{'*}\right)\right)\right] \\
& + \sum_{j^{'}=1}^{D^{'}}\sum_{t^{'}=1}^{T_{max}^{'}} \mathrm{E}_q \left[\log\left(p\left(\mu_{j^{'}t^{'}}|\varphi_{j^{'}t^{'}},\varsigma_{j^{'}t^{'}}\right)\right)\right] - \sum_{j^{'}=1}^{D^{'}}\sum_{t^{'}=1}^{T_{max}^{'}} \mathrm{E}_q \left[\log\left(q\left(\mu_{j^{'}t^{'}}|\varphi_{j^{'}t^{'}}^*,\varsigma_{j^{'}t^{'}}^*\right)\right)\right] \\
& + \sum_{j^{'}=1}^{D^{'}}\sum_{t^{'}=1}^{T_{max}^{'}} \mathrm{E}_q \left[\log\left(p\left(\delta_{j^{'}t^{'}}|\varpi_{j^{'}t^{'}},\chi_{j^{'}t^{'}}\right)\right)\right] - \sum_{j^{'}=1}^{D^{'}}\sum_{t^{'}=1}^{T_{max}^{'}} \mathrm{E}_q \left[\log\left(q\left(\delta_{j^{'}t^{'}}|\varpi_{j^{'}t^{'}}^*,\chi_{j^{'}t^{'}}^*\right)\right)\right] \\
& + \sum_{j^{'}=1}^{D^{'}} \mathrm{E}_q \left[\log\left(p\left(\mu_{j^{'}}^{'}|\varphi_{j^{'}}^{'},\varsigma_{j^{'}}^{'}\right)\right)\right] - \sum_{j^{'}=1}^{D^{'}} \mathrm{E}_q \left[\log\left(q\left(\mu_{j^{'}}^{'}|\varphi_{j^{'}}^{'*},\varsigma_{j^{'}}^{'*}\right)\right)\right] \\
& + \sum_{j^{'}=1}^{D^{'}} \mathrm{E}_q \left[\log\left(p\left(\delta_{j^{'}}^{'}|\varpi_{j^{'}}^{'},\chi_{j^{'}}^{'}\right)\right)\right] - \sum_{j^{'}=1}^{D^{'}} \mathrm{E}_q \left[\log\left(q\left(\delta_{j^{'}}^{'}|\varpi_{j^{'}}^{'*},\chi_{j^{'}}^{'*}\right)\right)\right]
\end{aligned}
\tag{1}
$$

To compute the variational expectations $E_q[.]$ in equation (1), we use the properties of exponential family distribution. If variational distributions for $q(\alpha_{jt}|\lambda_{jt})$ and $q(\beta_j|\iota_j)$ are Dirichlet distributions, then the exponential family representations are given by

$$
\begin{aligned}
q(\boldsymbol{\alpha}_t|\boldsymbol{\lambda}_t) &= \exp\left[\left(\sum_{j=1}^{D}\left(\lambda_j^t-1\right)\log\left(\alpha_j^t\right)\right)+\log\Gamma\left(\textstyle\sum_{j=1}^{D}\lambda_j^t\right)-\textstyle\sum_{j=1}^{D}\left(\log\Gamma\left(\lambda_j^t\right)\right)\right] \\
q(\boldsymbol{\beta}|\boldsymbol{\iota}) &= \exp\left[\left(\sum_{j=1}^{D}\left(\iota_j-1\right)\log\left(\beta_j\right)\right)+\log\Gamma\left(\textstyle\sum_{j=1}^{D}\iota_j\right)-\textstyle\sum_{j=1}^{D}\left(\log\Gamma\left(\iota_j\right)\right)\right]
\end{aligned}
$$

So the natural parameters and sufficient statistics of the Dirichlet distributions for $\alpha$ and $\beta$ are $\eta_{\alpha_{jt}}=\lambda_{jt}-1$, $T(\alpha_{jt})=\log(\alpha_{jt})$ and $\eta_{\beta_j}=\iota_j-1$, $T(\beta_j)=\log(\beta_j)$, respectively. The approximation expectations are

$$
\mathrm{E}[\alpha_{jt}] = \frac{\lambda_{jt}}{\sum_{j^{'}=1}^{D}\lambda_{tj^{'}}} \qquad \mathrm{E}[\beta_j] = \frac{\iota_j}{\sum_{j^{'}=1}^{S}\iota_{j^{'}}}
$$

$$
\mathrm{E}[\log(\alpha_{jt})] = \psi(\lambda_{jt}) - \psi\left(\sum_{j^{'}=1}^{D}\lambda_{kj^{'}}\right) \qquad \mathrm{E}[\log(\beta_j)] = \psi(\iota_j) - \psi\left(\sum_{j^{'}=1}^{S}\iota_{j^{'}}\right)
$$

where $\psi(.)$ is the digamma function.

Similarity, if variational distributions for $q\left(\mu_{j't'}|\varphi_{j't'},\varsigma_{j't'}\right)$ and $q\left(\delta_{j't'}|\varpi_{j't'},\chi_{j't'}\right)$ are norm and Wishart distributions, the approximation expectations are

$$\mathrm{E}\left[\mu_{j't'}\right]=\varphi_{j't'} \quad \begin{array}{l}\mathrm{E}\left[\delta_{j't'}\right]=\varpi_{j't'}\chi_{j't'}\\ \mathrm{E}\left[\log\left(\delta_{j't'}\right)\right]=\psi\left(\frac{\chi_{j't'}}{2}\right)D'log2\log\left(\varpi_{j't'}\right)\end{array}$$

where $\psi\left(.\right)$ is the digamma function.

### 3 Stochastic Optimization of the Variational Parameters for global clustering

With a truncated stick-breaking representation of the hierarchical Dirichlet process (HDP) [1], variational factors for the stick lengths $q\left(\Psi_k'|\vartheta_k,\vartheta_k'\right)$ are Beta distributions and the assignment variable $W_i^k=\mathrm{I}\left[W_i=k\right]$ for the $i^{th}$ sample allocation is governed by a multinomial distribution indexed by a variational parameter $g_{ik}$. The computation of the approximate expectation for the parameters of truncated stick-breaking representation has been considered carefully by [2]. The results of variational expectation for both variables are as follows:

$$\begin{array}{l}q\left(W_i=k\right)=g_i^k\\ q\left(W_i>k\right)=\sum\limits_{k'=k+1}^{K_{\max}}g_i^{k'}\\ \mathrm{E}_q\left[\log\left(\Psi_k'\right)\right]=\psi\left(\vartheta_k\right)-\psi\left(\vartheta_k+\vartheta_k'\right)\\ \mathrm{E}_q\left[\log\left(1-\Psi_k'\right)\right]=\psi\left(\vartheta_k'\right)-\psi\left(\vartheta_k+\vartheta_k'\right)\end{array}$$

Following the principles of the variational inference [1, 2], we consider the derivation of the update equation of variational parameters of $W_i^k$ for only one variable by fixing the others' distribution. The optimal coordinate updates are derived for these variational parameters. The log of the optimized factor to the posterior distribution of $W_i^k$ is

$$\log Q^*\left(W\right)=E_{Z,\Phi,\Psi,\epsilon,\epsilon',\alpha,\beta,\mu,\delta}\left[\log p\left(W,Z,\Phi,\Psi,\epsilon,\epsilon',\alpha,\beta,\mu,\delta\right)\right]+const=\sum_{i=1}^{N}\sum_{k=1}^{K}W_i^k\log\left(g_i^k\right)+const$$

$$\begin{array}{l}\log g_i^k=\\ \sum\limits_{t=1}^{T_{max}}\sum\limits_{j=1}^{D}\mathrm{E}_Q\left[\Phi_{1ij}\right]\mathrm{E}_Q\left[Z_{1kt}\right]\left(\mathrm{E}_Q\left[log\left(\frac{\Gamma\left(\sum\limits_{j=1}^{D}\alpha_{tj}\right)}{\Gamma\left(\sum\limits_{j=1}^{S}X_{1ij}+\sum\limits_{j=1}^{S}\alpha_{tj}\right)}\right)\right]+\sum\limits_{j=1}^{D}\mathrm{E}_Q\left[log\left(\frac{\Gamma(X_{1ij}+\alpha_{tj})}{\Gamma(\alpha_{tj})}\right)\right]+const\right)+\\ \sum\limits_{t'=1}^{T'_{max}}\sum\limits_{j'=1}^{D'}\mathrm{E}_Q\left[\Phi_{2ij'}\right]\mathrm{E}_Q\left[Z_{2kt'}\right]\left[\frac{1}{2}\mathrm{E}_Q\left[log(\delta_{t'j'})\right]+\left(\frac{-1}{2}\left(X_{2ij'}-\mathrm{E}_Q\left[\mu_{t'j'}\right]\right)^T\mathrm{E}_Q\left[\delta_{t'j'}\right]\left(X_{2ij'}-\mathrm{E}_Q\left[\mu_{t'j'}\right]\right)\right)\right]\\ +\mathrm{E}_Q\left[log(\Psi_k')\right]+\sum_{k'=1}^{k-1}\mathrm{E}_Q\left[log(1-\Psi_{k'}')\right]\end{array}\tag{2}$$

The expected logarithm of the functions $\mathrm{E}_Q\left[log\left(\frac{\Gamma(\sum_{j=1}^{D}\alpha_{tj})}{\Gamma(\sum_{j=1}^{D}X_{1ij}+\sum_{j=1}^{D}\alpha_{tj})}\right)\right]$ and $\mathrm{E}_Q\left[log\left(\frac{\Gamma(X_{1ij}+\alpha_{tj})}{\Gamma(\alpha_{tj})}\right)\right]$ in equation (2) have not the closed form. Thus, the calculation of these equations are analytically intractable. We need a closed-form expression to use the standard form of the variational inference or the stochastic variational inference [2]. In order to overcome this problem, we adopt the Taylor solutions in our previous paper [3]. The variational parameters are updated based on first-order Taylor expansion as follows:

$$\begin{array}{l}log\left(\frac{\Gamma(X_{1ij}+\alpha_{tj})}{\Gamma(\alpha_{tj})}\right)\geq log\left(\frac{\Gamma(X_{1ij}+\overline{\alpha_{tj}})}{\Gamma(\overline{\alpha_{tj}})}\right)+\frac{\partial F(\alpha_{tj})}{\partial\alpha_{tj}}\frac{\partial\alpha_{tj}}{\partial log(\alpha_{tj})}|_{\alpha_{tj}=\overline{\alpha_{tj}}}\left(log(\alpha_{tj})-log(\overline{\alpha_{tj}})\right)\\ \mathrm{E}_Q\left[log\left(\frac{\Gamma(X_{1ij}+\alpha_{tj})}{\Gamma(\alpha_{tj})}\right)\right]\geq log\left(\frac{\Gamma(X_{1ij}+\overline{\alpha_{tj}})}{\Gamma(\overline{\alpha_{tj}})}\right)+\overline{\alpha_{tj}}\left[\psi\left(\overline{\alpha_{tj}}+X_{1ij}\right)-\psi\left(\overline{\alpha_{tj}}\right)\right]\left(\mathrm{E}_Q\left[log(\alpha_{tj})\right]-log(\overline{\alpha_{tj}})\right)\\ \geq log\left(\frac{\Gamma(X_{1ij}+\overline{\alpha_{tj}})}{\Gamma(\overline{\alpha_{tj}})}\right)+\overline{\alpha_{tj}}\left[\psi\left(\overline{\alpha_{tj}}+X_{1ij}\right)-\psi\left(\overline{\alpha_{tj}}\right)\right]\left(\psi(\lambda_{kj}^*)-\psi\left(\sum_{j=1}^{D}\lambda_{tj}^*\right)-log(\overline{\alpha_{tj}})\right)\end{array}$$

$$\begin{array}{l}\mathrm{E}_Q\left[log\left(\frac{\Gamma(\sum_{j=1}^{D}\alpha_{tj})}{\Gamma(\sum_{j=1}^{D}X_{1ij}+\sum_{j=1}^{D}\alpha_{tj})}\right)\right]\geq log\left(\frac{\Gamma(\sum_{j=1}^{D}\overline{\alpha_{tj}})}{\Gamma\left(\sum_{j=1}^{D}X_{1ij}+\sum\limits_{j=1}^{D}\overline{\alpha_{tj}}\right)}\right)\\ +\sum\limits_{j=1}^{D}\left[\psi\left(\sum_{j=1}^{D}\overline{\alpha_{tj}}\right)-\psi\left(\sum_{j=1}^{D}X_{1ij}+\sum_{j=1}^{D}\overline{\alpha_{tj}}\right)\right]\overline{\alpha_{tj}}\left(\psi(\lambda_{tj}^*)-\psi\left(\sum_{j=1}^{D}\lambda_{tj}^*\right)-log(\overline{\alpha_{tj}})\right)\end{array}$$

Based on this approach, the variational parameters of $W_{ik}$ are updated as follow:

$$\log g_i^k = \sum_{t=1}^{T_{max}} \sum_{j=1}^{D} f_{ij} l_{1tk} log \left( \frac{\Gamma\left(\sum_{j=1}^{D} \overline{\alpha_{tj}}\right)}{\Gamma\left(\sum_{j=1}^{D} X_{1ij} + \sum_{j=1}^{D} \overline{\alpha_{tj}}\right)} \right)$$
$$+ \sum_{t=1}^{T_{max}} \sum_{j=1}^{D} f_{ij} l_{1tk} \left[ \psi\left(\sum_{j=1}^{D} \overline{\alpha_{tj}}\right) - \psi\left(\sum_{j=1}^{D} X_{1ij} + \sum_{j=1}^{D} \overline{\alpha_{tj}}\right) \right] \overline{\alpha_{tj}} \left( \psi(\lambda_{tj}^*) - \psi\left(\sum_{j=1}^{D} \lambda_{tj}^*\right) - log(\overline{\alpha_{tj}}) \right)$$
$$+ \sum_{t=1}^{T_{max}} \sum_{j=1}^{D} f_{ij} l_{1tk} \left[ log\left(\frac{\Gamma(X_{1ij} + \overline{\alpha_{tj}})}{\Gamma(\overline{\alpha_{tj}})}\right) + \overline{\alpha_{tj}} \left[\psi\left(\overline{\alpha_{tj}} + X_{1ij}\right) - \psi\left(\overline{\alpha_{tj}}\right)\right] \left( \psi(\lambda_{tj}^*) - \psi\left(\sum_{j=1}^{D} \lambda_{tj}^*\right) - log(\overline{\alpha_{tj}}) \right) \right]$$
$$+ \sum_{t'=1}^{T'_{max}} \sum_{j'=1}^{D'} f'_{ij'} l_{2kt'} \left( \frac{1}{2} \left[ \psi\left( \frac{\chi_{t'j'}^* + 1 - j'}{2} \right) + D' log2 + log|\varpi_{t'j'}^*| \right] + const \right)$$
$$+ \sum_{t'=1}^{T'_{max}} \sum_{j'=1}^{D'} f'_{ij'} l_{2kt'} \left( \frac{-1}{2} \left[ \chi_{t'j'}^* \left(X_{2ij'} - \varphi_{t'j'}^*\right) \varpi_{t'j'}^* \left(X_{2ij} - \varphi_{t'j'}^*\right)^T + \frac{D'}{\varsigma_{t'j'}^*} \right] \right)$$
$$+ \psi(\vartheta_k) - \psi(\vartheta_k + \vartheta'_k) + \sum_{k'=1}^{k-1} \psi(\vartheta'_{k'}) - \psi(\vartheta_{k'} + \vartheta'_{k'}) \quad (3)$$

Similarly, the optimized solution to the posterior distribution of unit-length sticks $\Psi'_{ik}$

$$\log Q^*\left(\Psi'_{ik}\right) = (1-1)\log\left(\Psi'_{ik}\right) + (\kappa - 1)\log\left(1 - \Psi'_{ik}\right)$$
$$+ \sum_{i=1}^{N} \mathrm{E}_Q\left[W_{ik}\right] \log\left(\Psi'_{ik}\right) + \sum_{i=1}^{N} \sum_{k'=k+1}^{K_{\max}} E_Q\left[W_{ik}\right] \log\left(1 - Psi'_{ik}\right) + const \quad (4)$$

which has the logarithmic form of the Beta distribution and the corresponding variational distributions of breaking proportions $q\left(\Psi'_{ik}\right)$ are considered to be the Beta distribution. Then, the standard conditions are satisfied for a closed form coordinate update for local parameters. Based on the stochastic natural gradient, the variational parameters of $\Psi'_{ik}$ are updated by computing variational expectations in equation (11) [4, 2],

$$(\vartheta_t)^{(t+1)} = \left(1 - \rho^{(t)}\right)(\vartheta_t)^{(t)} + \rho^{(t)} \left\{ 1 + \sum_{i=1}^{N} g_{ik} \right\}$$
$$\left(\vartheta'_t\right)^{(t+1)} = \left(1 - \rho^{(t)}\right)\left(\vartheta'_t\right)^{(t)} + \rho^{(t)} \left\{ \kappa + \sum_{i=1}^{N} \sum_{k'=k+1}^{K_{\max}} g_{ik} \right\}$$

### 4 Stochastic Optimization of the Variational Parameters for Dirichlet multinomial mixtures (DMM) Model

#### 4.1 Compute the variational function parameters for $Z_{1kt}$

Following the principles of the variational inference [4, 2], variational parameters of $Z_{1kt}$ is the local parameters, we consider the derivation of the update equation for only one variable by fixing the others' distribution. The optimal coordinate updates are derived for local parameters. The log of the optimized factor to the posterior distribution of $Z_{1kt}$ is

$$\log Q^*(Z_1) = E_{W,\Phi_1,\Pi_1,\epsilon,\alpha,\beta}\left[\log p\left(W, Z_1, \Phi_1, \Pi_1, \epsilon, \alpha, \beta\right)\right] + const = \sum_{k=1}^{K_{max}} \sum_{t=1}^{T_{max}} Z_{1kt} \log\left(l_{1kt}\right) + const$$

$$\log l_{1tk} = \sum_{i=1}^{N} \sum_{j=1}^{D} \mathrm{E}_Q\left[W_{ik}\right] \mathrm{E}_Q\left[\Phi_{1ij}\right] \left( \mathrm{E}_Q \left[ log \left( \frac{\Gamma\left(\sum_{j=1}^{D} \alpha_{tj}\right)}{\Gamma\left(\sum_{j=1}^{D} X_{1ij} + \sum_{j=1}^{D} \alpha_{tj}\right)} \right) \right] + \sum_{j=1}^{D} \mathrm{E}_Q \left[ log\left( \frac{\Gamma(X_{1ij} + \alpha_{tj})}{\Gamma(\alpha_{tj})} \right) \right] + const \right)$$
$$+ \mathrm{E}_Q\left[log(\Pi_{1t})\right] + \sum_{t'=1}^{t-1} \mathrm{E}_Q\left[log(1 - \Pi_{1t'})\right] \quad (5)$$

$$\mathrm{E}_Q\left[Z_{1kt}\right] = \exp\left\{\log\left(l_{1kt}\right)\right\}$$

The variational parameters of $Z_{ik}$ are updated as follow:

$$
\begin{aligned}
&\log l_{1kt} = \sum_{i=1}^{N}\sum_{j=1}^{D} g_{ik} f_{ij} log \left( \frac{\Gamma\left(\sum_{j=1}^{D}\overline{\alpha_{tj}}\right)}{\Gamma\left(\sum_{j=1}^{D} X_{1ij}+\sum_{j=1}^{D}\overline{\alpha_{tj}}\right)} \right) \\
&+\sum_{i=1}^{N}\sum_{j=1}^{D} g_{ik} f_{ij} \left[ \psi\left(\sum_{j=1}^{D}\overline{\alpha_{tj}}\right) - \psi\left(\sum_{j=1}^{D} X_{1ij} + \sum_{j=1}^{D}\overline{\alpha_{tj}}\right)\right] \overline{\alpha_{tj}} \left(\psi(\lambda_{tj}^{*}) - \psi\left(\sum_{j=1}^{D}\lambda_{tj}^{*}\right) - log(\overline{\alpha_{tj}})\right) \\
&+\sum_{i=1}^{N}\sum_{j=1}^{D} g_{ik} f_{ij} \left[ log\left(\frac{\Gamma(X_{1ij}+\overline{\alpha_{tj}})}{\Gamma(\overline{\alpha_{tj}})}\right) + \overline{\alpha_{tj}}\left[\psi\left(\overline{\alpha_{tj}} + X_{1ij}\right) - \psi\left(\overline{\alpha_{tj}}\right)\right] \left(\psi(\lambda_{tj}^{*}) - \psi\left(\sum_{j=1}^{D}\lambda_{tj}^{*}\right) - log(\overline{\alpha_{tj}})\right)\right] \\
&+\psi(\vartheta_t^{*}) - \psi(\vartheta_t^{*} + \vartheta_t^{*'}) + \sum_{t'=1}^{t-1}\psi(\vartheta_{t'}^{*'}) - \psi(\vartheta_{t'}^{*} + \vartheta_{t'}^{*'})
\end{aligned} \tag{6}
$$

#### 4.2 Compute the variational function parameters of indicator variables for microbiome selection

Similarity, the optimized solution to the posterior distribution of variable selection $\Phi_{1ij}$

$$
\log Q^{*}\left(\Phi_1\right) = E_{W,Z_1,\Pi_1,\epsilon,\alpha,\beta}\left[\log p\left(W, Z_1, \Phi_1, \Pi_1, \epsilon, \alpha, \beta\right)\right] + const = \sum_{i=1}^{N}\sum_{j=1}^{S}\Phi_{1ij}\log\left(f_{ij}\right) + const
$$

$$
\begin{aligned}
&\log f_{ij}^{\Phi_{1ij}} = \sum_{k=1}^{K_{max}}\sum_{t=1}^{T_{max}} g_{ik} l_{1tk} log \left( \frac{\Gamma\left(\sum_{j=1}^{D}\overline{\alpha_{tj}}\right)}{\Gamma\left(\sum_{j=1}^{D} X_{1ij}+\sum_{j=1}^{S}\overline{\alpha_{tj}}\right)} \right) \\
&+\sum_{k=1}^{K_{max}}\sum_{t=1}^{T_{max}} g_{ik} l_{1tk} \left[ \psi\left(\sum_{j=1}^{D}\overline{\alpha_{tj}}\right) - \psi\left(\sum_{j=1}^{D} X_{1ij} + \sum_{j=1}^{D}\overline{\alpha_{tj}}\right)\right] \overline{\alpha_{tj}} \left(\psi(\lambda_{tj}^{*}) - \psi\left(\sum_{j=1}^{D}\lambda_{tj}^{*}\right) - log(\overline{\alpha_{tj}})\right) \\
&+\sum_{k=1}^{K_{max}}\sum_{t=1}^{T_{max}} g_{ik} l_{1tk} \left[ log\left(\frac{\Gamma(X_{1ij}+\overline{\alpha_{tj}})}{\Gamma(\overline{\alpha_{tj}})}\right) + \overline{\alpha_{tj}}\left[\psi\left(\overline{\alpha_{tj}} + X_{1ij}\right) - \psi\left(\overline{\alpha_{tj}}\right)\right] \left(\psi(\lambda_{tj}^{*}) - \psi\left(\sum_{j=1}^{D}\lambda_{tj}^{*}\right) - log(\overline{\alpha_{tj}})\right)\right] \\
&+[\psi(\xi_1^{*}) - \psi(\xi_1^{*} + \xi_2^{*})]
\end{aligned} \tag{7}
$$

$$
\begin{aligned}
&\log f_{ij}^{1-\Phi_{1ij}} = log \left( \frac{\Gamma\left(\sum_{j=1}^{D}\overline{\beta_j}\right)}{\Gamma\left(\sum_{j=1}^{S} X_{1ij}+\sum_{j=1}^{D}\overline{\beta_j}\right)} \right) \\
&+\sum_{j=1}^{S} \left[ \psi\left(\sum_{j=1}^{D}\overline{\beta_j}\right) - \psi\left(\sum_{j=1}^{D} X_{1ij} + \sum_{j=1}^{D}\overline{\beta_j}\right)\right] \overline{\beta_j} \left(\psi(\iota_j^{*}) - \psi\left(\sum_{j=1}^{D}\iota_j^{*}\right) - log(\overline{\beta_j})\right) \\
&+log\left(\frac{\Gamma\left(X_{1ij}+\overline{\beta_j}\right)}{\Gamma(\overline{\beta_j})}\right) + \overline{\beta_j}\left[\psi\left(\overline{\beta_j} + X_{1ij}\right) - \psi\left(\overline{\beta_j}\right)\right] \left(\psi(\iota_j^{*}) - \psi\left(\sum_{j=1}^{D}\iota_j^{*}\right) - log(\overline{\beta_j})\right) \\
&+[\psi(\xi_2^{*}) - \psi(\xi_1^{*} + \xi_2^{*})]
\end{aligned} \tag{8}
$$

#### 4.3 Updating variational parameters of $\alpha$

Based on the principal framework of the stochastic variational inference [2], we consider parameters of $\alpha$ as global parameters. These parameters are updated by a stochastic gradient step and noisy estimations of the natural gradient of the variational objective with respect to $\alpha_{tj}$. Following the computational method of the natural gradient (Hoffman et al. 2013), we compute the natural gradient of equation (1) with respect to the global variational parameters of $\alpha_{tj}$.

We consider Dirichlet distribution as a prior distribution with concentration parameters $\lambda_{tj}$, so a conditional distribution of profile given the observation data has form of the Dirichlet distribution. We consider the representation of the exponential family,

$$
p\left(\alpha|W, Z_1, \Phi_1, \Pi_1, \epsilon, \beta\right) = h\left(\alpha\right)\exp\left(\eta\left(W, Z_1, \Phi_1, \Pi_1, \epsilon, \beta\right)^{T} t\left(\alpha\right) - a\left(\eta\left(W, Z_1, \Phi_1, \Pi_1, \epsilon, \beta\right)\right)\right)
$$

where: $h\left(.\right)$ is the base measure; $a\left(.\right)$ is the log-normalize; $\eta\left(.\right)$ is the natural parameter; $t\left(.\right)$ is the sufficient statistics.

As above assumption of the variational parameters, we set $q\left(\alpha|\lambda^{*}\right)$ to be Dirichlet distribution as the complete conditional distributions. So we get

$$
q\left(\alpha|\lambda^{*}\right) = h\left(\alpha\right)\exp\left(\left(\lambda^{*}\right)^{T} t\left(\pi\right) - a\left(\lambda^{*}\right)\right)
$$

We consider the lower bound for only $\alpha$,

$$\begin{aligned}
&\mathcal{L}(\alpha) = \mathrm{E}_Q\left[\log p\left(\alpha|W, Z_1, \Phi_1, \Pi_1, \epsilon, \beta\right)\right] - \mathrm{E}_Q\left[q\left(\alpha|\lambda^*\right)\right] \\
&= \mathrm{E}_Q\left[\log\left\{h(\alpha)\right\} + \eta\left(W, Z_1, \Phi_1, \Pi_1, \epsilon, \beta\right)^T t(\alpha) - a\left(\eta\left(W, Z_1, \Phi_1, \Pi_1, \epsilon, \beta\right)\right) - \log\left\{h(\alpha)\right\} - (\lambda^*)^T t(\alpha) + a(\lambda^*)\right] \\
&= \mathrm{E}_Q\left[\eta\left(W, Z_1, \Phi_1, \Pi_1, \epsilon, \beta\right)^T t(\alpha) - a\left(\eta\left(W, Z_1, \Phi_1, \Pi_1, \epsilon, \beta\right)\right) - (\lambda^*)^T t(\alpha) + a(\lambda^*)\right] \\
&= \mathrm{E}_Q\left[\eta\left(W, Z_1, \Phi_1, \Pi_1, \epsilon, \beta\right)\right]^T\left[\nabla_{\lambda^*}\left\{a(\lambda^*)\right\}\right] - a\left(\eta\left(W, Z_1, \Phi_1, \Pi_1, \epsilon, \beta\right)\right) - (\lambda^*)^T\left[\nabla_{\lambda^*}\left\{a(\lambda^*)\right\}\right] + a(\lambda^*)
\end{aligned}$$

where, the expected value of the sufficient statistics is the gradient of log normalizer $\mathrm{E}_Q\left[t(\alpha)\right] = \nabla_{\lambda^*}\left\{a(\lambda^*)\right\}$ If the classical principal of the gradient method for maximization is used directly to find a maximum of $\mathcal{L}(\alpha)$ based on taking step of size $\rho$ in direction of the gradient, the optimized results is

$$\lambda^*_{(t+1)} = \lambda^*_{(t)} + \rho\nabla_{(\lambda^*)}L(\alpha) = \lambda^*_{(t)} + \rho^{(t)}\left[\nabla^2_{(\lambda^*)}\left\{a(\lambda^*)\right\}\right]\left\{E_Q\left[\eta\left(W, Z_1, \Phi_1, \Pi_1, \epsilon, \beta\right)\right]^T - (\lambda^*)^T\right\}$$

Following [2], by premultiplying the gradient by the inverse Fisher information $\mathrm{G}(\lambda^*)$ and apply the stochastic natural gradient of the variational objective with respect to $\alpha$, we get

$$G(\lambda^*) = E_{\lambda^*}\left[\left(\nabla_{(\lambda^*)}\log q\left(\alpha|\lambda^*\right)\right)\left(\nabla_{(\lambda^*)}\log q\left(\alpha|\lambda^*\right)\right)^T\right] = \nabla^2_{(\lambda^*)}\left\{a(\lambda^*)\right\}$$

$$\widehat{\nabla_{(\lambda^*)}}L(\alpha) = \left\{G(\lambda^*)\right\}^{-1}\nabla_{(\lambda^*)}L(\alpha) = \left\{E_Q\left[\eta\left(W, Z_1, \Phi_1, \Pi_1, \epsilon, \beta\right)\right] - \lambda^*\right\}$$

$$\begin{aligned}
&\lambda^*_{(t+1)} = \lambda^*_{(t)} + \rho\widehat{\nabla_{(\lambda^*)}}L(\alpha) = \lambda^*_{(t)} + \rho\left\{E_Q\left[\eta\left(W, Z_1, \Phi_1, \Pi_1, \epsilon, \beta\right)\right] - \lambda^*_{(t)}\right\} \\
&= (1-\rho)\lambda^*_{(t)} + \rho\left\{E_Q\left[\eta\left(W, Z_1, \Phi_1, \Pi_1, \epsilon, \beta\right)\right]\right\}
\end{aligned} \tag{9}$$

Based traditional variational inference, we get the conditional distribution of $\alpha$

$$\begin{aligned}
&\log p\left(\alpha|W, Z_1, \Phi_1, \Pi_1, \epsilon, \beta\right) = \sum_{i=1}^{N}\sum_{j=1}^{D}\sum_{k=1}^{K_{max}}\sum_{t=1}^{T_{max}}\log\left(\frac{\Gamma(X_{1ij}+\alpha_{tj})}{\Gamma(\alpha_{tj})}\right) \times \log\left(\frac{\Gamma(X_{1ij}+\alpha_{tj})}{\Gamma(\alpha_{tj})}\right) \times \mathrm{E}_Q\left[W_{ik}\right]\mathrm{E}_Q\left[Z_{1kt}\right]\mathrm{E}_Q\left[\Phi_{1ij}\right] \\
&+ \log(\alpha_{tj})(\lambda_{tj} - 1) + const
\end{aligned}$$

$$\begin{aligned}
&\mathrm{E}_q\left[\eta\left(W, Z_1, \Phi_1, \Pi_1, \epsilon, \beta\right)\right] = \sum_{i=1}^{N}\sum_{k=1}^{K_{max}}\mathrm{E}_Q\left[W_{ik}\right]\mathrm{E}_Q\left[Z_{1kt}\right]\mathrm{E}_Q\left[\Phi_{1ij}\right]\overline{\alpha_{kj}}\times \\
&\left[\psi\left(\sum_{j=1}^{D}\overline{\alpha_{tj}}\right) - \psi\left(\sum_{j=1}^{D}X_{1ij} + \sum_{j=1}^{D}\overline{\alpha_{tj}}\right) + \psi\left(\overline{\alpha_{tj}} + X_{1ij}\right) - \psi\left(\overline{\alpha_{tj}}\right)\right]
\end{aligned} \tag{10}$$

By substituting equation (10) into equation (9), the final result of updated variational parameters is

$$\begin{aligned}
&\left(\lambda^*_{tj}\right)^{(t+1)} = \left(1-\rho^{(t)}\right)\left(\lambda^*_{tj}\right)^{(t)} \\
&+\rho^{(t)}\left\{\lambda_{tj} + \sum_{i=1}^{N}\sum_{k=1}^{K_{max}}g_{ik}l_{1kt}f_{ij}\overline{\alpha_{tj}}\left[\psi\left(\sum_{j=1}^{D}\overline{\alpha_{tj}}\right) - \psi\left(\sum_{j=1}^{D}X_{1ij} + \sum_{j=1}^{D}\overline{\alpha_{tj}}\right) + \psi\left(\overline{\alpha_{tj}} + X_{1ij}\right) - \psi\left(\overline{\alpha_{tj}}\right)\right]\right\}
\end{aligned}$$

#### 4.4 Updating variational parameters of $\beta$

Similarity, the stochastic natural gradient of the variational objective with respect to $\beta$

$$\begin{aligned}
&\left(\iota^*_j\right)^{(t+1)} = \left(1-\rho^{(t)}\right)\left(\iota^*_j\right)^{(t)} \\
&+\rho^{(t)}\left\{\iota_j + \sum_{i=1}^{N}\left[1-f_{ij}\right]\overline{\beta_j}\left(\left[\psi\left(\sum_{j=1}^{D}\overline{\beta_j}\right) - \psi\left(\sum_{j=1}^{D}X_{1ij} + \sum_{j=1}^{D}\overline{\beta_j}\right)\right] + \left[\psi\left(\overline{\beta_j} + X_{1ij}\right) - \psi\left(\overline{\beta_j}\right)\right]\right)\right\}
\end{aligned}$$

#### 4.5 Updating stick-breaking representation

Similarity, the optimized solution to the posterior distribution of unit length sticks $\Pi^{'}_{1t}$

$$\begin{aligned}
&\log Q^*\left(\Pi^{'}_{1t}\right) = (1-1)\log\left(\Pi^{'}_{1t}\right) + (\kappa^* - 1)\log\left(1 - \Pi^{'}_{1t}\right) \\
&+ \sum_{k=1}^{K_{max}}\mathrm{E}_Q\left[Z_{1tk}\right]\log\left(\Pi^{'}_{1t}\right) + \sum_{k=1}^{K_{max}}\sum_{t'=t+1}^{T_{\max}}E_Q\left[Z_{1kt'}\right]\log\left(1 - \Pi^{'}_{1t}\right) + const
\end{aligned} \tag{11}$$

which has the logarithmic form of the Beta distribution and the corresponding variational distributions of breaking proportions $q(\Pi_{1t})$ are considered to be the Beta distribution. Then, the standard conditions are satisfied for a closed form coordinate update for local parameters. Based on the stochastic natural gradient, the variational parameters of $\Pi_{1t}$ are updated by computing variational expectations in equation (11) [4, 2],

$$(\vartheta_t^*)^{(t+1)} = \left(1-\rho^{(t)}\right)(\vartheta_t^*)^{(t)} + \rho^{(t)}\left\{1+\sum_{k=1}^{K_{max}} l_{1kt}\right\}$$
$$\left(\vartheta_t^{*'}\right)^{(t+1)} = \left(1-\rho^{(t)}\right)\left(\vartheta_t^{*'}\right)^{(t)} + \rho^{(t)}\left\{\kappa^* + \sum_{k=1}^{K_{max}}\sum_{t'=t+1}^{T_{\max}} l_{1kt'}\right\}$$

#### 4.6 Updating variational parameters of $\epsilon$

Similarity, the stochastic natural gradient of the variational objective with respect to $\epsilon$

$$\left(\xi_{j1}^*\right)^{(t+1)} = \left(1-\rho^{(t)}\right)\left(\xi_{j1}^*\right)^{(t)} + \rho^{(t)}\left\{\xi_{j1} + \sum_{i=1}^{N} f_{ij}\right\}$$
$$\left(\xi_{j2}^*\right)^{(t+1)} = \left(1-\rho^{(t)}\right)\left(\xi_{j2}^*\right)^{(t)} + \rho^{(t)}\left\{\xi_{j2} + \sum_{i=1}^{N}\left(1-f_{ij}\right)\right\}$$

### 5 Stochastic Optimization of the Variational Parameters for Gaussian mixture model (GMM)

#### 5.1 Compute the variational function parameters for $Z_{2kt'}$

Following the principles of the variational inference [4, 2], variational parameters of $Z_{2kt'}$ is the local parameters, we consider the derivation of the update equation for only one variable by fixing the others' distribution. The optimal coordinate updates are derived for local parameters. The log of the optimized factor to the posterior distribution of $Z_{2kt'}$ is

$$\log Q^*\left(Z_2\right) = E_{W,\Phi_2,\Pi_2,\epsilon',\mu,\delta,\mu',\delta'}\left[\log p\left(W, Z_2, \Phi_2, \Pi_2, \epsilon', \mu, \delta, \mu', \delta'\right)\right] + const = \sum_{k=1}^{K_{max}}\sum_{t'=1}^{T'_{max}} Z_{2kt'}\log\left(l_{2kt'}\right) + const$$

$$\begin{aligned}
&\log l_{2kt'} = \\
&\sum_{i=1}^{N}\sum_{j'=1}^{D'} \mathrm{E}_Q\left[W_{ik}\right]\mathrm{E}_Q\left[\Phi_{2ij'}\right]\left[\tfrac{1}{2}\mathrm{E}_Q\left[log(\delta_{t'j'})\right] + \left(\tfrac{-1}{2}\left(X_{2ij} - \mathrm{E}_Q\left[\mu_{t'j'}\right]\right)^T \mathrm{E}_Q\left[\delta_{t'j'}\right]\left(X_{2ij} - \mathrm{E}_Q\left[\mu_{t'j'}\right]\right)\right)\right] \\
&+\mathrm{E}_Q\left[log(\Pi_{2t'})\right] + \sum_{t''=1}^{t'-1}\mathrm{E}_Q\left[log(1-\Pi_{2t''})\right]
\end{aligned} \tag{12}$$

$$\mathrm{E}_Q\left[Z_{2kt'}\right] = \exp\left\{\log\left(l_{2kt'}\right)\right\}$$

The variational parameters of $Z_{2kt'}$ are updated as follow:

$$\begin{aligned}
\log l_{2kt'} &= \sum_{i=1}^{N}\sum_{j'=1}^{D'} g_{ik}f'_{ij}\left(\tfrac{1}{2}\left[\psi\left(\frac{\chi^*_{t'j'}+1-j}{2}\right) + D'log2 + log|\varpi^*_{t'j'}|\right]\right) \\
&+\sum_{i=1}^{N}\sum_{j'=1}^{D'} g_{ik}f'_{ij'}\left(\tfrac{-1}{2}\left[\chi^*_{t'j'}\left(X_{2ij'} - \varphi^*_{t'j'}\right)\varpi^*_{t'j'}\left(X_{2ij'} - \varphi^*_{t'j'}\right)^T + \frac{D'}{\varsigma^*_{t'j'}}\right]\right) \\
&+\psi(\vartheta^+_{t'}) - \psi(\vartheta^+_{t'} + \vartheta^{+'}_{t'}) + \sum_{t''=1}^{t'-1}\psi(\vartheta^{+'}_{t''}) - \psi(\vartheta^+_{t''} + \vartheta^{+'}_{t''})
\end{aligned} \tag{13}$$

#### 5.2 Compute the variational function parameters of indicator variables for metabolome selection

Similarity, the optimized solution to the posterior distribution of variable selection $\Phi_{2ij'}$

$$\log Q^*\left(\Phi_2\right) = E_{W,Z_2,\Pi_2,\epsilon',\mu,\delta,\mu',\delta'}\left[\log p\left(W, Z_2, \Phi_2, \Pi_2, \epsilon', \mu, \delta, \mu', \delta'\right)\right] + const = \sum_{i=1}^{N}\sum_{j'=1}^{D'}\Phi_{2ij'}\log\left(f'_{ij'}\right) + const$$

$$\log f_{ij'}^{\Phi_{2ij'}} = \sum_{k=1}^{K_{max}} \sum_{t'=1}^{T'_{max}} g_{ik} l_{2kt'} \left( \frac{1}{2} \left[ \psi \left( \frac{\chi^*_{t'j'}+1-j}{2} \right) + D' log2 + log|\varpi^*_{t'j'}| \right] + const \right)$$
$$+ \sum_{i=1}^{N} \sum_{j'=1}^{D'} g_{ik} f'_{ij'} \left( \frac{-1}{2} \left[ \chi^*_{t'j'} \left( X_{2ij'} - \varphi^*_{t'j'} \right) \varpi^*_{t'j'} \left( X_{2ij'} - \varphi^*_{t'j'} \right)^T + \frac{D'}{\varsigma^*_{t'j'}} \right] \right) \tag{14}$$
$$+ \left[ \psi(\xi'^*_1) - \psi(\xi'^*_1 + \xi'^*_2) \right]$$

$$\log f_{ij'}^{1-\Phi_{2ij'}} = \left( \frac{1}{2} \left[ \psi \left( \frac{\chi'^*_{j'}+1-j'}{2} \right) + D' log2 + log|\varpi'^*_{j'}| \right] + const \right)$$
$$+ \left( \frac{-1}{2} \left[ \chi'^*_{j'} \left( X_{2ij'} - \varphi'^*_{j'} \right) \varpi'^*_{j'} \left( X_{2ij'} - \varphi'^*_{j'} \right)^T + \frac{D'}{\varsigma'^*_{j'}} \right] \right) \tag{15}$$
$$+ \left[ \psi(\xi'^*_2) - \psi(\xi'^*_1 + \xi'^*_2) \right]$$

#### 5.3 Updating stick-breaking representation

The optimized solution to the posterior distribution of unit length sticks $\Pi'_{2t'}$

$$\log Q^* \left( \Pi'_{2t'} \right) = (1-1) \log \left( \Pi'_{2t'} \right) + (\kappa^+ - 1) \log \left( 1 - \Pi'_{2t'} \right)$$
$$+ \sum_{k=1}^{K_{max}} \mathrm{E}_Q \left[ Z_{2kt'} \right] \log \left( \Pi'_{2t'} \right) + \sum_{k=1}^{K_{max}} \sum_{t''=t'+1}^{T'_{\max}} E_Q \left[ Z_{2kt''} \right] \log \left( 1 - \Pi'_{2t''} \right) + const \tag{16}$$

Based on the stochastic natural gradient, the variational parameters of $\Pi'_{2t}$ are updated by computing variational expectations in equation (16) [4, 2],

$$\left( \vartheta^+_{t'} \right)^{(t+1)} = \left( 1 - \rho^{(t)} \right) \left( \vartheta^+_{t'} \right)^{(t)} + \rho^{(t)} \left\{ 1 + \sum_{k=1}^{K_{max}} l_{2kt'} \right\}$$
$$\left( \vartheta^{+'}_{t'} \right)^{(t+1)} = \left( 1 - \rho^{(t)} \right) \left( \vartheta^{+'}_{t'} \right)^{(t)} + \rho^{(t)} \left\{ \kappa^+ + \sum_{k=1}^{K_{max}} \sum_{t''=t'+1}^{T'_{\max}} l_{2kt''} \right\}$$

#### 5.4 Updating variational parameters of $\epsilon'$

The stochastic natural gradient of the variational objective with respect to $\epsilon'$

$$\left( \xi'^*_{j'1} \right)^{(t+1)} = \left( 1 - \rho^{(t)} \right) \left( \xi'^*_{j'1} \right)^{(t)} + \rho^{(t)} \left\{ \xi'_{j'1} + \sum_{i=1}^{N} f'_{ij'} \right\}$$
$$\left( \xi'^*_{j'2} \right)^{(t+1)} = \left( 1 - \rho^{(t)} \right) \left( \xi'^*_{j'2} \right)^{(t)} + \rho^{(t)} \left\{ \xi'_{j'2} + \sum_{i=1}^{N} \left( 1 - f'_{ij'} \right) \right\}$$

#### 5.5 Updating variational parameters of $\mu$

The stochastic natural gradient of the variational objective with respect to $\mu$

$$\left( \varphi^*_{j't'} \right)^{(t+1)} = \left( 1 - \rho^{(t)} \right) \left( \varphi^*_{j't'} \right)^{(t)} + \rho^{(t)} \left\{ \frac{\varphi_{j't'} \varsigma_{j't'} + \sum_{i=1}^{N} \sum_{k=1}^{K_{max}} g_{ik} l_{2kt'} f'_{ij'} X_{2ij'}}{\varsigma_{j't'} + \sum_{i=1}^{N} \sum_{k=1}^{K_{max}} g_{ik} l_{2kt'} f'_{ij'}} \right\}$$
$$\left( \varsigma^*_{j't'} \right)^{(t+1)} = \left( 1 - \rho^{(t)} \right) \left( \varsigma^*_{j't'} \right)^{(t)} + \rho^{(t)} \left\{ \varsigma_{j't'} + \sum_{i=1}^{N} \sum_{k=1}^{K_{max}} g_{ik} l_{2kt'} f'_{ij'} \right\}$$

#### 5.6 Updating variational parameters of $\delta$

The stochastic natural gradient of the variational objective with respect to $\delta$

$$\left( \varpi^*_{j't'} \right)^{(t+1)} = \left( 1 - \rho^{(t)} \right) \left( \varpi^*_{j't'} \right)^{(t)} +$$
$$\rho^{(t)} \left\{ \varpi_{j't'} + \sum_{i=1}^{N} \sum_{k=1}^{K_{max}} g_{ik} l_{2kt'} f'_{ij'} \left( X_{2ij'} - \varphi^*_{j't'} \right) \left( X_{2ij'} - \varphi^*_{j't'} \right)^T + \frac{\varsigma_{j't'} \sum_{i=1}^{N} \sum_{k=1}^{K_{max}} g_{ik} l_{2kt'} f'_{ij'}}{\varsigma_{j't'} + \sum_{i=1}^{N} \sum_{k=1}^{K_{max}} g_{ik} l_{2kt'} f'_{ij'}} \left( \varphi^*_{j't'} - \varphi_{j't'} \right)^T \left( \varphi^*_{j't'} - \varphi_{j't'} \right) \right\}$$
$$\left( \chi^*_{j't'} \right)^{(t+1)} = \left( 1 - \rho^{(t)} \right) \left( \chi^*_{j't'} \right)^{(t)} + \rho^{(t)} \left\{ \chi_{j't'} + \sum_{i=1}^{N} \sum_{k=1}^{K_{max}} g_{ik} l_{2kt'} f'_{ij'} \right\}$$

##### 5.7 Updating variational parameters of $\mu^{'}$

The stochastic natural gradient of the variational objective with respect to $\mu^{'}$

$$\left(\varphi_{j'}^{'*}\right)^{(t+1)} = \left(1-\rho^{(t)}\right)\left(\varphi_{j'}^{'*}\right)^{(t)} + \rho^{(t)}\left\{\frac{\varphi_{j'}^{'}\varsigma_{j'}^{'}+\sum_{i=1}^{N} f_{ij'}^{'}X_{2ij'}}{\varsigma_{j'}^{'}+\sum_{i=1}^{N} f_{ij'}^{'}}\right\}$$

$$\left(\varsigma_{j'}^{'*}\right)^{(t+1)} = \left(1-\rho^{(t)}\right)\left(\varsigma_{j'}^{'*}\right)^{(t)} + \rho^{(t)}\left\{\varsigma_{j'}^{'}+\sum_{i=1}^{N} f_{ij'}^{'}\right\}$$

##### 5.8 Updating variational parameters of $\delta^{'}$

The stochastic natural gradient of the variational objective with respect to $\delta^{'}$

$$\left(\varpi_{j'}^{'*}\right)^{(t+1)} = \left(1-\rho^{(t)}\right)\left(\varpi_{j'}^{'*}\right)^{(t)} +$$

$$\rho^{(t)}\left\{\varpi_{j'}^{'}+\sum_{i=1}^{N} f_{ij'}^{'}\left(X_{2ij'}-\varphi_{j'}^{'*}\right)\left(X_{2ij'}-\varphi_{j'}^{'*}\right)^{T}+\frac{\varsigma_{j'}^{'}\sum_{i=1}^{N} f_{ij'}^{'}}{\varsigma_{j'}^{'}+\sum_{i=1}^{N} f_{ij'}^{'}}\left(\varphi_{j'}^{'*}-\varphi_{j'}^{'}\right)^{T}\left(\varphi_{j'}^{'*}-\varphi_{j'}^{'}\right)\right\}$$

$$\left(\chi_{j'}^{'*}\right)^{(t+1)} = \left(1-\rho^{(t)}\right)\left(\chi_{j'}^{'*}\right)^{(t)} + \rho^{(t)}\left\{\chi_{j'}^{'}+\sum_{i=1}^{N} f_{ij'}^{'}\right\}$$

### 6 Soybean rhizosphere microbiome and metabolome

#### 6.1 Field experiment

The accessions and experimental fields were identical to those used by Toda et al. and Sakurai et al [5, 6]. A diverse panel of 198 soybean accessions registered in the National Agriculture and Food Research Organization Genbank (https://www.gene.affrc.go.jp/) was used. These 198 accessions mainly consist of Japanese and global soybean minicore collections [7, 8]. The field trial was conducted in 2019 in an experimental field with sandy soil at the Arid Land Research Center, Tottori University (35°32’ N, 134°12’ E, 14 m above sea level). Each plot consisted of four plants, and the distances between the two rows, two plots, and two individuals were 50, 80, and 20 cm, respectively. Sowing was performed at the beginning of July, followed by thinning after two weeks. Fertilizers (13, 6.0, 20, 11, and 7.0 g m-2 of N, P, K, Mg, and Ca, respectively) were applied to the field before sowing. White mulch sheets (Dupont, Wilmington, DE, USA) were laid to prevent rainwater infiltration and to control soil conditions with artificial irrigation. Watering tubes were installed under the sheets to irrigate fields. Two watering treatments, non-watered, and well-watered, were used to evaluate the influence of drought and control conditions. The watering treatment was started after thinning and two weeks after sowing. Artificial irrigation was applied at a flow rate of 1.1L / h m for 5 h daily (7:00–9:00, 12:00–14:00, and 16:00–17:00). Root sampling was conducted at the beginning of September. Root sampling was performed from on the 3rd individual. A root of an individual was collected and stored at -80 degrees Celsius until metabolome and microbiome analysis.

#### 6.2 Metabolome analysis

Metabolome analysis of root samples was performed according to Sawada et al., 2009 [9] and Uchida et al., 2020 [10]. The collected samples were freeze-dried and powdered using a mixer (Shake Master NEO BMS-M10N21, Bio Medical Science, Tokyo, Japan). The powdered sample was weighed to 4 mg, 1 mL of extraction solvent (80% methanol containing 0.1% formic acid, 8.4 nM lidcain, and 210 nM 10-camphoresulfonic acid as internal standards) was added, and sonication was performed for 10 min. After centrifugation (10,000 rpm for 5 min), the collected supernatant was used as the extract. To 25 $\mu$L of the extract, 75 $\mu$L of the extraction solvent was added. Twenty-five $\mu$L of the diluted extract was dried with N2 gas, and 250 $\mu$L of deionized water was added, dissolved by shaking, and used as the test solution. The final concentrations of the internal standards in the test solutions of the root samples were 0.84 nM for lidcain and 21 nM for 10-camphoresulfonic acid. After filtration through a MultiScreen 386 well-plate filter (Merck Millipore, Billerica, MA, USA), the test solution was subjected to metabolome analysis using a Shimadzu LCMS-8050 system coupled to a Nexera X2 UPLC system (Shimadzu, Kyoto, Japan). A Waters HSS T3 column (1.0 × 50 mm, 1.8 $\mu$m) was used, and the column oven temperature was kept at 30 °C. The mobile phase consisted of 0.1% (v/v) formic acid/water (A) and 0.1% (v/v) formic acid/acetonitrile (B) with gradient elution of 0.1 to 9% (B) at 0.25–0.4 min, 9 to 17% (B) at 0.4–0.8 min, 17 to 99.9% (B) at 0.8–1.9 min, 99.9 to 99.9% (B) at 1.9–2.1 min, 99.9 to 0.1% (B) at 2.1–2.11 min, and 0.1% (B) at 2.11–2.7 min. The flow rate of the mobile phase was 0.24 mL/min and a sample injection volume of

1 $\mu$L. The mass spectrometer was operated in the positive (+) and negative (-) ESI modes. Parameters for the interface were set as follows: nebulizing gas flow 3 mL/min, heating gas flow 10 L/min, drying gas flow 10 L/min, interface temperature 300 °C, dilution line temperature 250 °C, heat block temperature 400 °C, collision-induced dissociation gas 270 kPA and capillary voltage 4 kV (+); -3.5kv (-). The mass spectrometer was operated in the scheduled multiple reaction monitoring (MRM) mode for MS/MS measurements. The MRM transitions of each compound are listed in Supplementary Table 1. Raw data (lcd file) were converted into Abf files using the Reifycs Abf (Analysis base file) Converter (https://www.reifycs.com/AbfConverter/). Peak areas of the LC–QqQ-MS data were calculated using MRMPROBS (Tsugawa et al. 2013, http://prime.psc.riken.jp/compms/mrmprobs/main.html) [11].

#### 6.3 Microbiome analysis

##### DNA extraction

Bacterial DNA was extracted from the root and rhizosphere soils, as previously described [12, 13]. Briefly, 100 mg of the powdered sample was dispensed into a 1.5 mL tube and chilled in liquid nitrogen. Subsequently, 200 $\mu$L of Lysis/binding buffer (LBB) containing 1 M LiCl (Cat. #L7026-500ML, Sigma-Aldrich, St. Louis, MO, USA), 100 mM Tris-HCl (Cat. #318-90225, Wako Pure Chemical Corporation, Osaka, Japan), 1% SDS (Cat. #313-90275, Wako Pure Chemical Corporation), 10 mM EDTA pH 8.0 (Cat. #311-90075, Wako Pure Chemical Corporation), Antifoam A (Cat. #A5633-25G, Sigma-Aldrich), 5 mM dithiothreitol (Cat. #048-29224, Wako Pure Chemical Corporation), 11.2 M 3-Mercapto-1,2-propanediol (Cat. #139-16452, Wako Pure Chemical Corporation) and DNase/RNase-free H2O (Cat. #10977015, Thermo Fisher Scientific, Waltham, MA, USA) [14] were added to the tubes, mixed by vortexing, incubated at room temperature (24 °C) for 5 min. The tube of each sample was centrifuged at 13,000 rpm for 10 min at RT and 50 $\mu$L of supernatant was mixed with an equal amount of AMPure XP beads (Cat. #A63881, Beckman Coulter, Inc.) in new tube to clean and size selection of DNA. The mixture was vortexed, incubated for 5 min at RT and then placed on a magnetic station for 5 min. After removing the supernatant, the magnetic beads were washed twice with 200 $\mu$L of 80% ethanol. Finally, the DNA was eluted from the magnetic beads using 15 $\mu$L of 10 mM Tris-HCl (pH 7.5).

##### Bacterial 16S rRNA gene sequencing

Libraries were prepared by modifying a previously described two-step PCR amplification protocol [12, 14]. Briefly, the DNA library was constructed using primers (515f: 5'- TCG TCG GCA GCG TCA GAT GTG TAT AAG AGA CAG - [3-6-mer Ns] - GTG YCA GCM GCC GCG GTA A -3', 806rB: 5'- GTC TCG TGG GCT CGG AGA TGT GTA TAA GAG ACA G - [3-6-mer Ns] - GGA CTA CNV GGG TWT CTA AT -3') for amplifying the V4 region of the bacterial 16S rRNA [15, 16]. The V4 region of 16S rRNA was amplified in a 10 $\mu$L reaction containing 1 $\mu$L of 10-fold diluted sample, 2× KAPA HiFi HotStart ReadyMix (Cat. #07958935001, KAPA Biosystems Inc., USA), 0.2 $\mu$M forward and reverse primers, and 1 $\mu$M blocking primers (mPNA and pPNA; PNA BIO, Inc., Newbury Park, CA, USA) in a thermal cycler as follows: 1 cycle for 3 min at 95 °C, 35 cycles for 20 sec at 98 °C, 10 sec at 78 °C, 30 sec at 50 °C, 90 sec at 72 °C, and a final extension for 5 min at 72 °C (ramp rate = 1 °C/s).

The first PCR product was purified using ExoSAP-IT Express (Cat #75001.1.EA; Thermo Fisher Scientific) to remove excess primer and dNTPs. The PCR product (5 $\mu$L) was mixed with 2 $\mu$L of ExoSAP-IT Express and incubated at 37 °C for 4 min, followed by inactivation at 80 °C for 1 min.

The second PCR was performed using the following index primers: forward primer (5'- AAT GAT ACG GCG ACC ACC GAG ATC TAC AC - [8-mer index] - TCG TCG GCA GCG TC -3') and reverse primer (5'- CAA GCA GAA GAC GGC ATA CGA GAT - [8-mer index] - GTC TCG TGG GCT CGG -3') [17]. The second PCR reaction was performed in a 10 $\mu$L reaction containing 0.8 $\mu$L of the purified first amplification, 2× KAPA HiFi HotStart ReadyMix, 1 $\mu$M forward and reverse primers, and 1 $\mu$M blocking primers (mPNA and pPNA) in a thermal cycler as follows: 1 cycle for 3 min at 95 °C, 8 cycles for 20 sec at 98 °C, 10 sec at 78 °C, 30 sec at 55 °C, 90 sec at 72 °C, and a final extension for 5 min at 72 °C (ramp rate = 1 °C/s). For the second PCR amplification, each sample was mixed with AMPure XP beads, vortexed and left at room temperature for 5 min. The samples were then placed in a magnetic station for 5 min. After removing the supernatant, the magnetic beads were washed twice with 200 $\mu$L of 80% ethanol. The DNA libraries were eluted from the magnetic beads with 15 $\mu$L of 10 mM Tris-HCl (pH 7.5) and quantified using Quant-iT PicoGreen dsDNA reagent (Cat. #P7581, Invitrogen) and a microplate photometer (Infinite 200 PRO M Nano+, TECAN Japan Co., Ltd., Kanagawa, Japan). DNA libraries were diluted and pooled with equimolar concentrations of each library. The pooled libraries were quantified by qPCR using the NEBNext Library Quant Kit for Illumina (Cat. #E7630L, New England BioLabs, Ipswich, MA, USA) and diluted to 3 nM. Denatured libraries were spiked with 20% PhiX Control v3 (Cat. #FC-110-3001, Illumina, San Diego, CA, USA) and sequenced on an Illumina MiSeq platform using a 2× 300-bp MiSeq Reagent Kit v3 (Illumina).

##### 16S V4 rRNA data preprocessing

Raw paired-end reads were processed using the Quantitative Insight Into Microbial Ecology 2 program (QIIME2, ver. 2020.6.0). Raw FASTQ files were imported into QIIME2, sequencing primers were trimmed using the Cutadapt plugin. For ASV-based analysis, the primer-free sequencing reads were analyzed using the DADA2 pipeline [18] to truncate forward and reverse reads (length of 240 for forward reads and 180 for reverse reads), filter, denoise, merge reads, and detect and remove chimeras. The taxonomy of all ASVs was annotated using SILVA (ver. 138) reference database [19, 20]. Sequences derived from Archaea, Eukaryota, mitochondria, or chloroplasts were removed.
